# Novel Predictive Spatial Biomarker in Non-Small Cell Lung Carcinoma: The Diversity of Niches Unlocking Treatment Sensitivity (DONUTS)

**DOI:** 10.1101/2025.08.13.665980

**Authors:** Tricia R. Cottrell, Jeffrey S. Roskes, Michael Fotheringham, Emily Cohen, Boyang Zhang, Logan L Engle, Daphne Wang, Elizabeth Will, Joel C. Sunshine, Daniel Jimenez-Sanchez, Zhen Zeng, Justina X. Caushi, Jiajia Zhang, Nina M. D’Amiano, Julie S. Deutsch, Sonali Uttam, Katie Pirie, Darah Vlaminck, Michelle Mataj, Eman Radwan, Alexa Fiorante, Nicole Espinosa, Teodora Popa, Aleksandra Ogurtsova, Sigfredo Soto-Diaz, Margaret Eminizer, Samuel Tabrisky, Andrew Jorquera, Jonathan Skidmore, Dmitry Medvedev, Jamie E. Chaft, Julie R. Brahmer, Michael Conroy, Joshua E. Reuss, Ludmila Danilova, Hongkai Ji, Patrick M. Forde, Drew M. Pardoll, Kellie N. Smith, Benjamin F. Green, Alexander S. Szalay, Janis M. Taube

## Abstract

Probabilistic spatial modelling techniques developed on large-scale tumor-immune Atlases (∼35M individually mapped cells; 50,000 high power fields) were used to characterize predictive features of treatment-responsive lung cancer. We identified CD8+FoxP3+ cell density as a robust pre-treatment biomarker for outcomes across disease stages and therapy types. In parallel, single-cell RNAseq studies of CD8+FoxP3+ T-cells revealed an activated, early effector phenotype, substantiating an anti-tumor role, and contrasting with CD4+FoxP3+ T-regulatory cells. A spatial biomarker was developed using an empirical probabilistic model to define the immediate cell neighbors or niche surrounding CD8+FoxP3+ cells and proximity to the tumor-stromal boundary. The resultant ‘Diversity of Niches Unlocking Treatment Sensitivity (DONUTS)’ are more prevalent than the CD8+FoxP3+ cells themselves, mitigating sampling error in small biopsies. Further, the DONUTS only require four markers, are additive to PD-L1, and associate with tertiary lymphoid structure counts. Taken together, the DONUTS represent a next-generation predictive biomarker poised for clinical implementation.

**HIGHLIGHTS:**

- Large-scale tumor-immune Atlases drive robust computational biomarker development
- CD8+FoxP3+ cells are anti-tumor T-cells and predict response to therapy
- The niches or spatial ‘donuts’ around CD8+FoxP3+ cells boost biomarker performance
- CD8+FoxP3+ donuts are hallmarks of a larger immune organization that includes TLS

## INTRODUCTION

Immune checkpoint blockade (ICB) of the programmed cell death protein 1 (PD-1) receptor or its ligand, programmed cell death ligand 1 (PD-L1), is now the standard first-line therapy for patients with many advanced cancers, including non-small cell lung cancer (NSCLC). In patients with resectable NSCLC, treatment with neoadjuvant or perioperative anti-PD-(L)1-based regimens improves event-free survival.^1–4^ In both settings, pre-treatment tumor cell PD-L1 expression has been shown to enrich for subsequent response to anti-PD-(L)1. Well-recognized limitations include limited sensitivity and specificity, which are largely attributable to the detection of a single parameter within the tumor microenvironment (TME). As such, it is an incomplete representation of the multiplicity of cells contributing to response as well as the multiparameter PD-1/PD-L1 receptor-ligand interactions targeted by these agents.^5,6^ Moreover, there have been numerous reports describing the association of tertiary lymphoid structures (TLS) with response to ICB, supporting the role of the larger tumor ecosystem in orchestrating an effective anti-tumor immune response.^7,8^ However, TLS are best detected in large resection specimens, and therefore are not typically evident in small, pre-treatment core biopsies.^9,10^ More effective pre-treatment biomarkers remain highly sought after.

Multiplex immunofluorescence (mIF) is an emerging technology that allows visualization of several immune markers simultaneously, and has been shown to out-perform other biomarkers such as PD-L1 immunohistochemistry (IHC), interferon-gamma gene signature, and mutational density for predicting response to anti-PD-(L)1.^11^ Because mIF preserves the tissue architecture of the TME, it supports characterization of the local niches and larger ecosystem underpinning the efficacy of immune checkpoint blockade.^12–14^ The increasingly complex spatial organization patterns revealed using these emerging technologies require distinct statistical approaches for quantification, ranging from first-order statistics (e.g., densities of individual cell lineages) to third- and higher-order statistics (e.g., spatial relationships across three or more elements in the TME).^15,16^ While these approaches are well developed in fields like astronomy and geography, many of these strategies have not yet been adapted to capture relevant biological information embedded in cellular maps of tumors.

Most TME spatial profiling studies to date narrowly distill highly complex spatial datasets into simplified variables to support analytic feasibility and ease of interpretation (e.g., characterizing pairwise heterotypic cellular interactions and/or homotypic cell clustering). These limited features are extracted from data-rich cellular maps and systematically tested for associations with patient outcomes. Despite the data lost with such approaches, these initial efforts have captured the non-random distribution of immune cells in the TME and identified novel biomarker candidates.^17–19^ The continued evolution of multi-parameter approaches to TME characterization will require (1) overcoming the loss of higher-order spatial information, (2) substantiation of putative discovered biology using orthogonal methods, (3) validation in multiple, independent cohorts, and (4) subsequent biomarker assay development to optimize signal and clinical feasibility. Addressing these limitations is expected to produce robust spatial biomarkers for clinical translation.

Defining features of current clinical pathology practices include the use of whole slide specimens and a limited number of IHC or immunofluorescent labels, which support tractable, cost-effective workflows and timely results reporting. Whole slides also have the benefit of providing templates for studying the higher-order immunoarchitecture and associated spatial biology that is not possible to define using tissue microarrays (TMA) or select high power fields (HPFs). The breadth of the data and sheer number of cells acquired in whole slide strategies allow for rare cells to be identified with statistical certainty. For example, we previously identified CD8+FoxP3+ cells as being strongly associated with response to anti-PD-1-based regimens in patients with melanoma using whole slide mapping, despite the fact that they constitute only 2-3% of all CD8+ T-cells in the TME.^20^ That study was made possible by generating and querying maps using the AstroPath platform, an end-to-end workflow inspired by large astronomical surveys.^21^ AstroPath was designed to generate rigorously calibrated tumor-immune maps with highly-accurate single cell resolution.

Here, we present pre- and on-treatment mIF tumor and immune AstroPath Atlases of NSCLC whole slide maps, with accompanying single cell RNA sequencing (scRNAseq) data. The Atlases include both squamous and non-squamous tumor specimens collected across multiple independent cohorts and treatment settings – including neoadjuvant and advanced disease, chemotherapy and anti-PD-1 immunotherapy, and those treated as a part of clinical trials and as standard of care. Using these Atlases, we extend our previous efforts and show that the CD8+FoxP3+ cells are also a strong predictor of therapeutic response in NSCLC, and use scRNAseq to characterize them as early, effector tumor-specific T-cells. We also define their biological niche with regard to their immediate contact neighbors (*i.e.,* the ‘donut’ of cells surrounding the CD8+FoxP3+ cells), as well as their higher-order spatial organization relative to the tumor-stromal boundary. We use this information to develop a biologically-informed, multidimensional spatial model for predicting therapeutic response that requires only 4 markers, is additive to PD-L1, and is a surrogate for the presence of TLS. The resultant spatial biomarker, the Diversity of Niches Unlocking Treatment Sensitivity (DONUTS), is associated with patient outcomes across tumor stage and treatment settings. These mIF whole slide Atlases are comprised of 35M coordinately-plotted cells assembled from ∼50,000 HPFs, providing a robust, resource dataset sufficient to support the development of stable computational models, facilitating biomarker discovery poised for clinical implementation.

## RESULTS

### Cohort summary

We studied 75 pre-treatment biopsy specimens from four cohorts of patients with NSCLC treated with systemic therapy, **Figure 1A**. Cohort 1 (n=25) was from the first neoadjuvant clinical trial of anti-PD-1 (NCT02259621)^22^; Cohort 2 (n=14) was from a landmark trial of advanced NSCLC receiving second-line anti-PD-1 (NCT01673867)^23^; Cohort 3 (n=20) received second-line standard of care anti-PD-1; and Cohort 4 (n=16) was from a landmark trial of advanced NSCLC receiving second-line chemotherapy (docetaxel) following progression on first-line platinum doublet chemotherapy (NCT01673867)^23^. We tested all pre-treatment biopsies from patients meeting inclusion criteria who had tissue available for study (see **Figure S1** for CONSORT diagrams). We also characterized 35 on-treatment, definitive resection specimens from Cohort 1.

**Figure 1.**
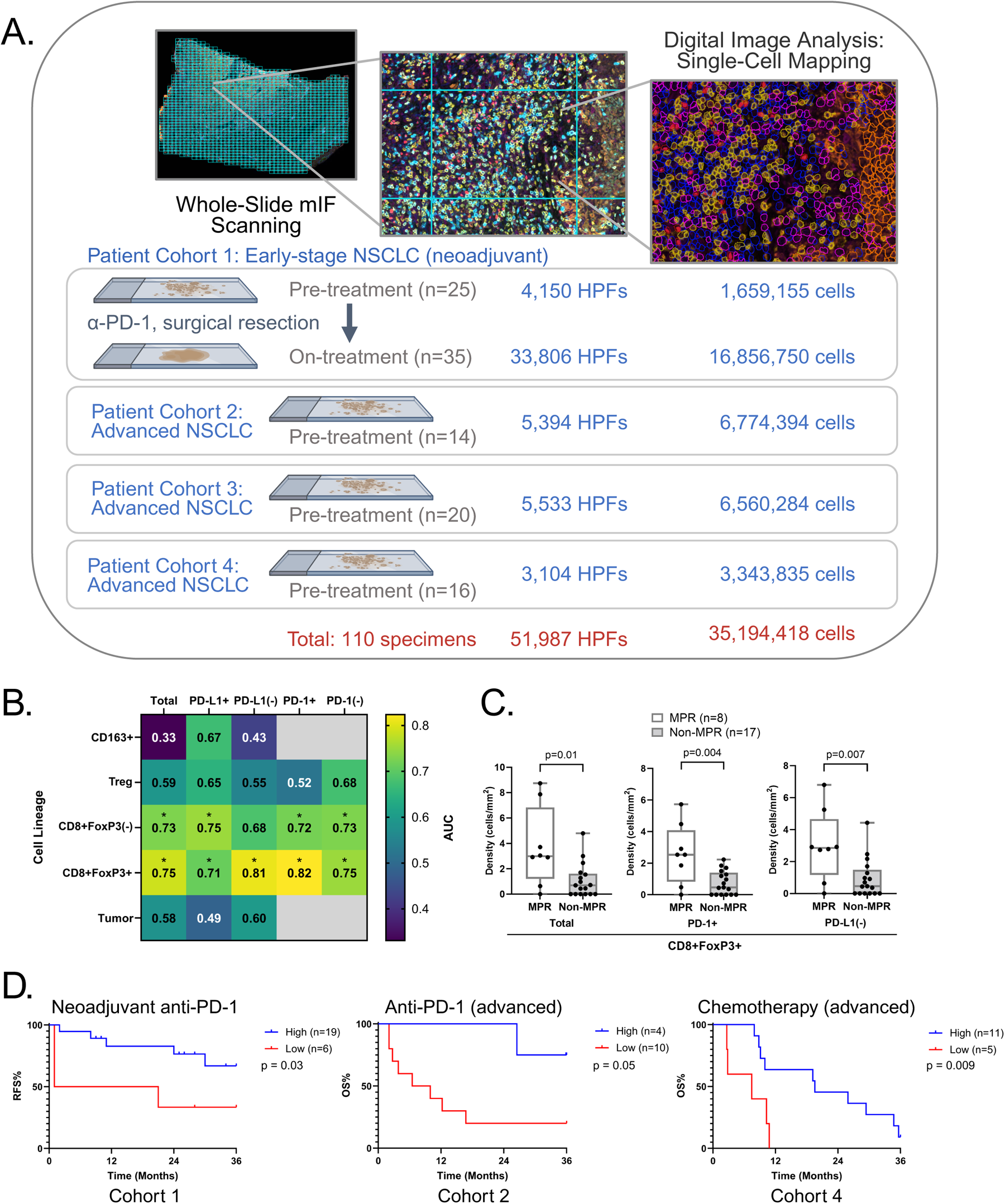
CD8+FoxP3+ cell densities predict patient outcomes to therapy in both the neoadjuvant and advanced NSCLC treatment settings. (A) Study schematic summarizing patient cohorts, tumor tissue specimens, high powered fields (HPFs), and single cells mapped using the AstroPath platform. (B) Heatmap of pre-treatment cell density AUC values for prediction of major pathologic response to neoadjuvant anti-PD-1-based therapy. (C) Boxplots showing the comparison of total pre-treatment CD8+FoxP3+ cell densities between MPR and non-MPR (regardless of PD-1 and PD-L1 status) as well as the select subsets by PD-(L)1 status with the highest AUCs. Boxplots of the other cell lineage densities (CD163+ macrophages, CD8+ T-cells, conventional T-regs, tumor) as they relate to MPR are shown in Figure S2B-E. (D) Pre-treatment CD8+FoxP3+ cell densities stratify (*left*) RFS in patients with early-stage NSCLC treated with neoadjuvant anti-PD-1-based therapy (Cohort 1) as well as (*middle*) OS in patients with advanced, pre-treated NSCLC receiving anti-PD-1 therapy (Cohort 2) and (*right*) chemotherapy (Cohort 4). Cohort 3 did not have survival data available. OS for Cohort 1 and PFS for Cohorts 2 and 4 are shown in Figure S2F-H. (* p < 0.05) MPR: Major pathologic response; RFS: recurrence free survival; OS: overall survival.

Clinicopathologic data was gathered for the patients included in each cohort and included age, sex, NSCLC, histologic subtype, treatment regimen received, RECIST 1.1 objective radiographic response to therapy and/or pathologic response to therapy, recurrence free survival (RFS), progression free survival (PFS), and overall survival (OS), when available, **Tables S1-3**.^24^

The resultant tumor-immune Atlas is composed of ∼50,000 HPFs and more than 35 million individually mapped cells as well as the corresponding clinical outcomes data. This resource can be accessed using instructions provided in **Table S4**.

### CD8+FoxP3+ T-cells predict survival in patients receiving anti-PD-1 or chemotherapy

A validated six-marker multiplex immunofluoresence (mIF) assay (PD-1, PD-L1, CD8, FoxP3, CD163 and pan-CK) was performed on pre-treatment biopsies from Cohort 1, **Figure S2A, Table S5**. The assay was designed to detect individual cell types, including CD163+ macrophages, CD8+ T cells, CD8(-)FoxP3+ regulatory T cells (Treg cells), CD8+FoxP3+ T-cells, tumor cells, and “other” (negative for these markers), as well as PD-1 and PD-L1 expression on each of these cell lineages. The association of the pre-treatment cell densities for each lineage with major pathologic response (MPR, ≤10% residual viable tumor) after neoadjuvant therapy was assessed by calculating the area under the curve (AUC) for the receiver operator characteristic (ROC) curve (**Figure 1B**). Among all lineages evaluated, the highest AUC value (0.75) was observed for the relatively rare CD8+FoxP3+ cell population (3.4% of all CD8+ T-cells and 0.11% of all cells within the pre-treatment TME). These cells were significantly enriched in pre-treatment biopsies from patients who went on to demonstrate a major pathologic response to neoadjuvant anti-PD-1 (**Figure 1C**). Refined CD8+FoxP3+ cellular subsets based on expression of PD-1 and PD-L1 further strengthened the association with MPR, *e.g.*, CD8+FoxP3+PD-1+ (AUC 0.82) (**Figure 1B-C**). CD8+ cell densities also showed an association with outcomes, though it was not as strong as for the CD8+FoxP3+ cells (**Figure S2B-E**). The recurrence-free survival (RFS) among patients with CD8+FoxP3+ T-cells was also improved (**Figure 1D**). Median overall survival (OS) has not yet been reached, though those with CD8+FoxP3+ T-cells have shown a durable improvement in OS (89% vs. 50% 2-year OS, p=0.05, respectively, **Figure S2F)**.

To extend our understanding of CD8+FoxP3+ T-cells across tumor stage, histologic subtypes, and treatment settings, we performed the mIF assay on pre-treatment biopsies from two cohorts of patients with advanced NSCLC treated with anti-PD-1. Cohort 2 included patients with adenocarcinomas and Cohort 3 included patients with a mixture of adenocarcinomas and squamous cell carcinomas. In Cohort 2, patients with high CD8+FoxP3+ cell densities demonstrated significantly improved overall survival (OS) (100% vs. 27% 2-year OS, respectively p<0.0001; median OS: not reached vs. 8.2 months, **Figure 1D**) and a trend towards improved PFS **(Figure S2G)**. In Cohort 3, pre-treatment CD8+FoxP3+ cell densities predicted radiographic response to therapy (AUC 0.73). PFS and OS data were not available for this cohort.

Chemotherapy has historically been regarded as immunosuppressive. However, emerging data suggests some traditional chemotherapeutic agents may alter the tumor microenvironment to stimulate antitumor immunity.^25^ We tested this in patients who received docetaxel chemotherapy in the second-line after progression on platinum doublet chemotherapy (Cohort 4). In keeping with our hypothesis, we showed that higher CD8+FoxP3+ T-cell densities in biopsy specimens taken prior to the second-line chemotherapy is associated with improved PFS and OS (median 9.0 vs. 2.2 months, p<0.001; and median 19.6 vs. 7.5 months, p=0.009, respectively, **Figure 1D, Figure S2H).**

### The CD8+FoxP3+ T-cell signature is additive to PD-L1 IHC

PD-L1 IHC is the current clinical standard for predicting response to anti-PD-(L)1-based therapy. We combined the assessment of CD8+FoxP3+ density with PD-L1 tumor cell expression using a 50% threshold. We found that in both early-stage (neoadjuvant, Cohort 1) and advanced NSCLC (Cohort 2), the combination of these markers improved OS stratification compared to PD-L1 alone (p<0.0001 and p=0.01, respectively, **Figure 2A**), indicating non-redundant contributions. As noted previously, long-term survival data was not available for Cohort 3, and PD-L1 did not associate with patient outcomes following chemotherapy in the parent clinical trial for Cohort 4.^23^

**Figure 2.**
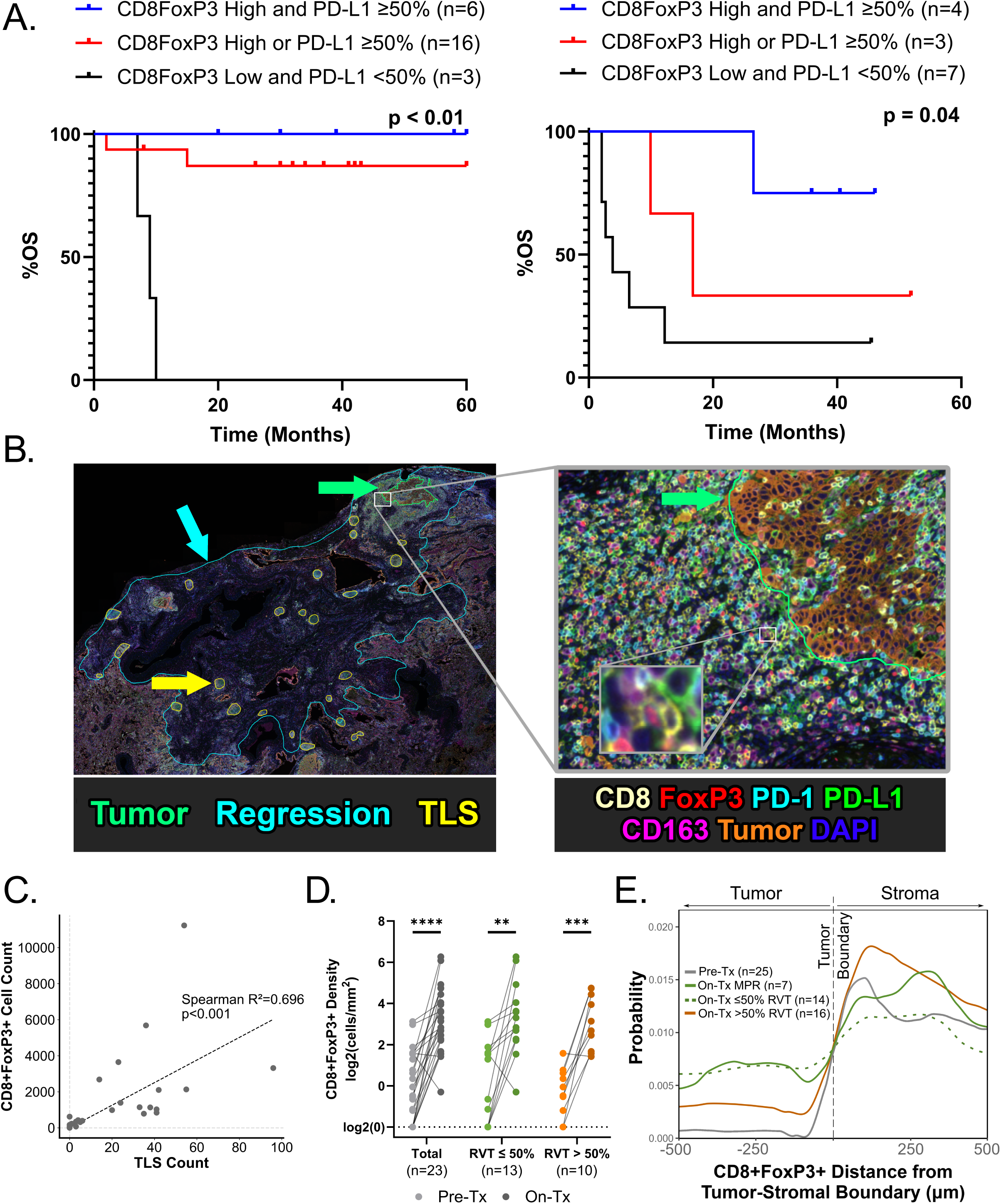
CD8+FoxP3+ cell density is additive to PD-L1, indicative of TLS, and increases during anti-PD-1 treatment response. (**A**) Kaplan-Meier plots of (*left*) OS in Cohort 1 and (*right*) OS in Cohort 2 demonstrate enhanced patient stratification by combined assessment of pre-treatment tumor PD-L1 expression and CD8+FoxP3+ T-cell densities. **(B)** (*Left*) An on-treatment mIF TME map with annotations outlining residual tumor (green), regression bed (i.e. where tumor used to be prior to treatment, blue) of a patient who experienced a major pathologic response (≤10% residual viable tumor). TLS (yellow) are readily identified in the TME, and many are remote from where the residual tumor is located. (*Right*) A high magnification image of the tumor-stromal boundary illustrates a dense immune cell population immediately adjacent and infiltrating the residual tumor cells (orange), including a mixture of CD8+ cytotoxic T cells (yellow), CD8+FoxP3+ cells (yellow and red, see inset), FoxP3+ regulatory T cells (red), and CD163+ macrophages (purple). **(C)** Correlation of CD8+FoxP3+ cell counts with TLS in on-treatment resection specimens from Cohort 1. **(D)** CD8+FoxP3+ cell densities are significantly increased following neoadjuvant therapy regardless of pathologic response status. Shown here are those with a partial pathologic response (≤50% RVT) vs. those >50% RVT, with a trend towards higher densities in patients with a deeper pathologic response (p=0.087). **(E)** Probability distribution of spatial localization of CD8+FoxP3+ cells relative to the tumor-stromal boundary. Relative to pre-treatment tumors (gray), CD8+FoxP3+ cells infiltrate the tumor epithelium in on-treatment specimens with at least partial pathologic response (≤50% RVT, green), while these cells are largely confined to the peritumoral stroma in on-treatment tumors with >50% RVT (orange). (** p < 0.005, *** p < 0.0005). TLS: tertiary lymphoid structures; RVT: residual viable tumor

### CD8+FoxP3+ T-cell density positively correlates with the number of tertiary lymphoid structures (TLS)

Neoadjuvant treatment offers a unique opportunity to extensively profile the TME in on-treatment tumor specimens. We generated whole-slide mIF maps of the on-treatment tumor resections from Cohort 1 and were able to readily identify TLS, in addition to other TME features (**Figure 2B**). We and others have previously reported that TLS are readily identified in on-treatment NSCLC resections and are associated with response to anti-PD-(L)1 therapy.^7–9^ We identified a positive correlation between the densities of CD8+FoxP3+ T-cells within the tumor bed and the number of TLS in on-treatment specimens (p<0.001) (**Figure 2C**).

### Pre vs. on-treatment dynamics and spatial distribution of CD8+FoxP3+ T-cells

We used paired pre- and on-treatment specimens from Cohort 1 to characterize the dynamics and spatial distribution of CD8+FoxP3+ T-cells under therapeutic pressure. The densities of CD8+FoxP3+ T-cells significantly increased in the tumor bed following exposure to anti-PD-1 (**Figure 2D**). When comparing patients who experienced a MPR vs. non-MPR, there was a trend toward higher CD8+FoxP3+ densities in the on-treatment tumors in those with MPR (median 11 cells/mm^2^ vs. 6 cells/mm^2^, respectively). Evaluation of the spatial distribution of CD8+FoxP3+ T cells showed a 3-fold increase into tumors with evidence of at least partial pathologic response after treatment (Wilcoxon rank-sum test, p=0.008), suggesting a role in anti-tumor immunity (**Figure 2E**).

### On-treatment CD8+FoxP3+ T-cell scRNAseq profiling substantiates a role in anti-tumor immunity

Collectively, the dynamics of CD8+FoxP3+ T-cells in response to anti-PD-1, including cellular proliferation, tumor infiltration, and correlation with TLS, suggest these cells may be a hallmark of an effective anti-tumor immune response. To define the functional profile of CD8+FoxP3+ T-cells, we reanalyzed a previously published single-cell RNA sequencing (scRNA-seq) dataset of tumor infiltrating lymphocytes isolated from (n=15) patients from Cohort 1 treated with neoadjuvant anti-PD-1.^26^ Uniform manifold approximation and projection (UMAP) dimensionality reduction defined 11 FoxP3+ T-cell clusters (**Figure 3A**). In contrast to conventional, suppressive CD4+FoxP3+ Tregs, cluster-defining genes for the CD8+FoxP3+ T-cell subset were pro-inflammatory and cytotoxic (including CCL5, NKG7, CTSW, and granzymes B, A and K, adjusted p-value ≤ 0.05) **(Figure S3).**

**Figure 3:**
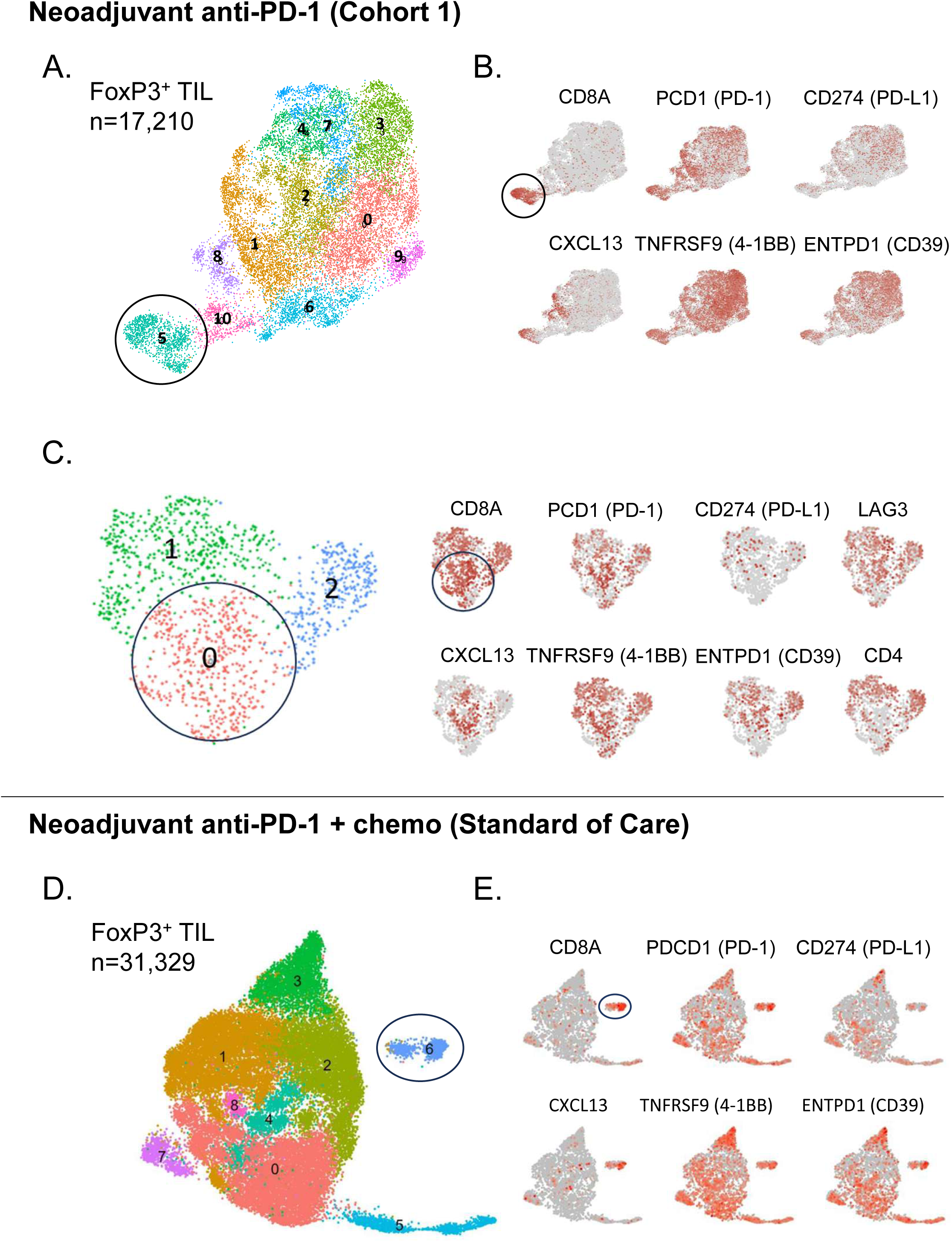
Tumor infiltrating CD8+FoxP3+ T-cells have an activated, cytotoxic phenotype in patients treated with anti-PD-1+/-chemotherapy. **(A)** Single-cell RNA sequencing-derived UMAP of the expression profiles of the FoxP3+ TIL, resulting in 11 unique immune cell subsets (Cohort 1, see also **Figure S3)**. **(B)** UMAP overlay of expression of select genes indicates antigen-specific activation of the CD8+FoxP3+ subset **(C)** Subclustering of CD8+FoxP3+ cells in cluster 5 further highlights the CXCL13 expression in CD8+CD4-FoxP3+ T-cells. (**D-E**) UMAP clustering and expression overlays of scRNAseq data from FoxP3+ TIL from patients receiving neoadjuvant combination chemo-immunotherapy as standard of care.^30^ A similar transcriptional program consistent with antigen-specific activation is identified in the CD8+ cluster of FoxP3+ TIL (circled, see also **Figure S4**). TIL: tumor infiltrating lymphocytes.

Deeper interrogation of the CD8+FoxP3+ cells by subclustering highlighted CXCL13 expression, one of the most selective genes for tumor neoantigen-specific CD8 T-cells (**Figure 3B-C**.^26–28^ Gene set enrichment analysis was then performed to compare this population to resting CD8 cells, specifically stem-like memory cells (Table S6). The CD8+FoxP3+ T-cells were enriched for an activation signature (HLA class II, cytotoxic granules, and perforins). Further, top genes associated with suppressive CD4+ Tregs^26^ were not identified in the top 30 enriched genes (*e.g*., OX40, GITR, IL1R2, LAYN, CD25, EBI3). These findings were then replicated in an independent scRNA-seq dataset obtained from n=222 patients who received standard of care neoadjuvant anti-PD-1+chemotherapy (**Figure 3D-E**, Figure S4).^29^ Taken together, these findings indicate CD8+FoxP3+ T-cells are highly-potent early, effector T-cells and explain their strong positive association with immunotherapy treatment response.

### Pre-treatment CD8+FoxP3+ T-cells localize to a distinct cellular niche

The AstroPath tumor-immune maps were used to characterize the contact neighbors and spatial distribution of CD8+FoxP3+ T-cells. The CD8+FoxP3+ *niche* was defined by the ‘donut’ or ring of cells surrounding a CD8+FoxP3+ cell (**Figure 4A**) as well as its distance from the tumor-stromal boundary (**Figure 4B**, see **Table 1** for definitions of spatial arrangement terminology used herein). In pre-treatment specimens, CD8+FoxP3+ T-cells were localized to the peritumoral stroma, consistent with prior findings in melanoma.^20^ In all four NSCLC cohorts, the proportion of CD8+ cytotoxic T-cells was enriched >3.5-fold among contact neighbors of CD8+FoxP3+ T-cells compared to the background TME (**Figure S5A**, 37.3% vs 6.3%, p=0.01; 52.0% vs. 12.2%, p<0.001; 32.4% vs. 6.7%, p<0.001; and 45.0% vs. 12.3%, p<0.001 in Cohorts 1, 2, 3, and 4, respectively). Conversely, the proportion of neighboring tumor cells was reduced >4-fold relative to their abundance in the background TME, while other lineages were not consistently altered (**Figure S5B-E**).

**Figure 4.**
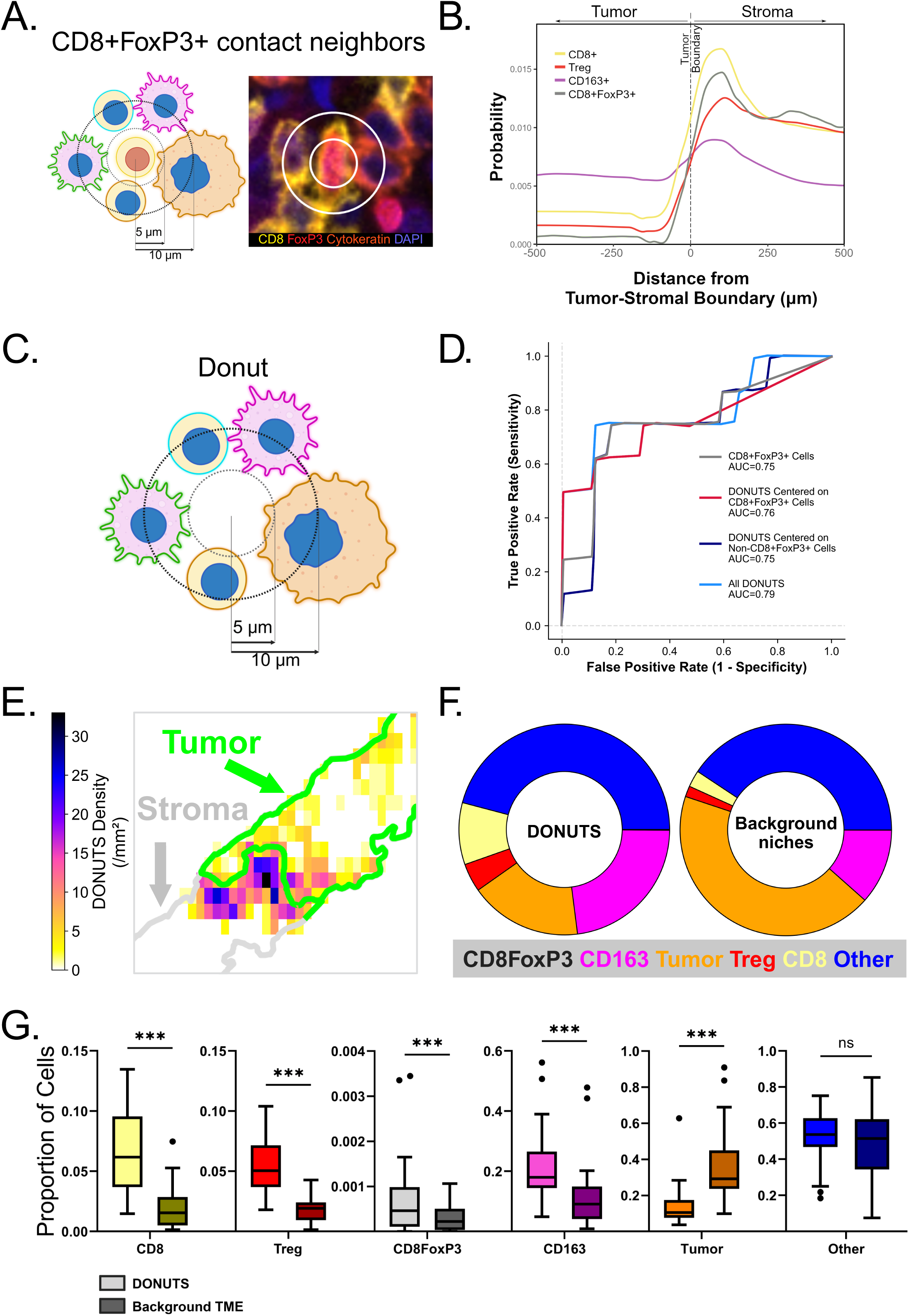
CD8+FoxP3+ T-cells localize to a T-cell enriched immunoactive niche near the tumor-stromal boundary. **(A)** Schematic and photomicrograph showing the ring or donut-like arrangement of immediate contact neighbors of CD8+FoxP3+ cells**. (B)** Probability distribution of spatial localization of each cell lineage relative to the tumor-stromal boundary in pre-treatment biopsies from Cohort 1 shows that CD8+FoxP3+ cells, like CD8+ cells and Tregs, are most concentrated in the stroma in close proximity to the tumor-stromal interface. (**C)** Schematic of an immunoactive niche derived from the CD8+FoxP3+ cell niche, but without the requirement of a central cell, resulting in a ring or a donut-shaped cellular arrangement. The probability model defined the subset of these niches predictive of treatment response, termed ‘DONUTS’ – the Diversity Of Niches Unlocking Treatment Sensitivity. (**D**) ROC curves for predicting MPR in Cohort 1 based on pre-treatment densities of (i) CD8+FoxP3+ cells, (ii) DONUTS centered on CD8+FoxP3+ cells, (iii) DONUTS centered on non-CD8+FoxP3+ cells, and (iv) all DONUTS. (**E**) Heat map showing the localization of DONUTS to the tumor-stromal boundary in a representative pre-treatment NSCLC biopsy. **(F)** Pie charts comparing the cellular composition of the DONUTS relative to non-predictive background niches. **(G)** Boxplots comparing the proportions of each cell lineage in the background TME vs. the DONUTS, which mirror the composition observed around actual CD8+FoxP3+ T-cells (**Figure S5**). (*** p < 0.0005; ns: not significant).

**Table 1.**
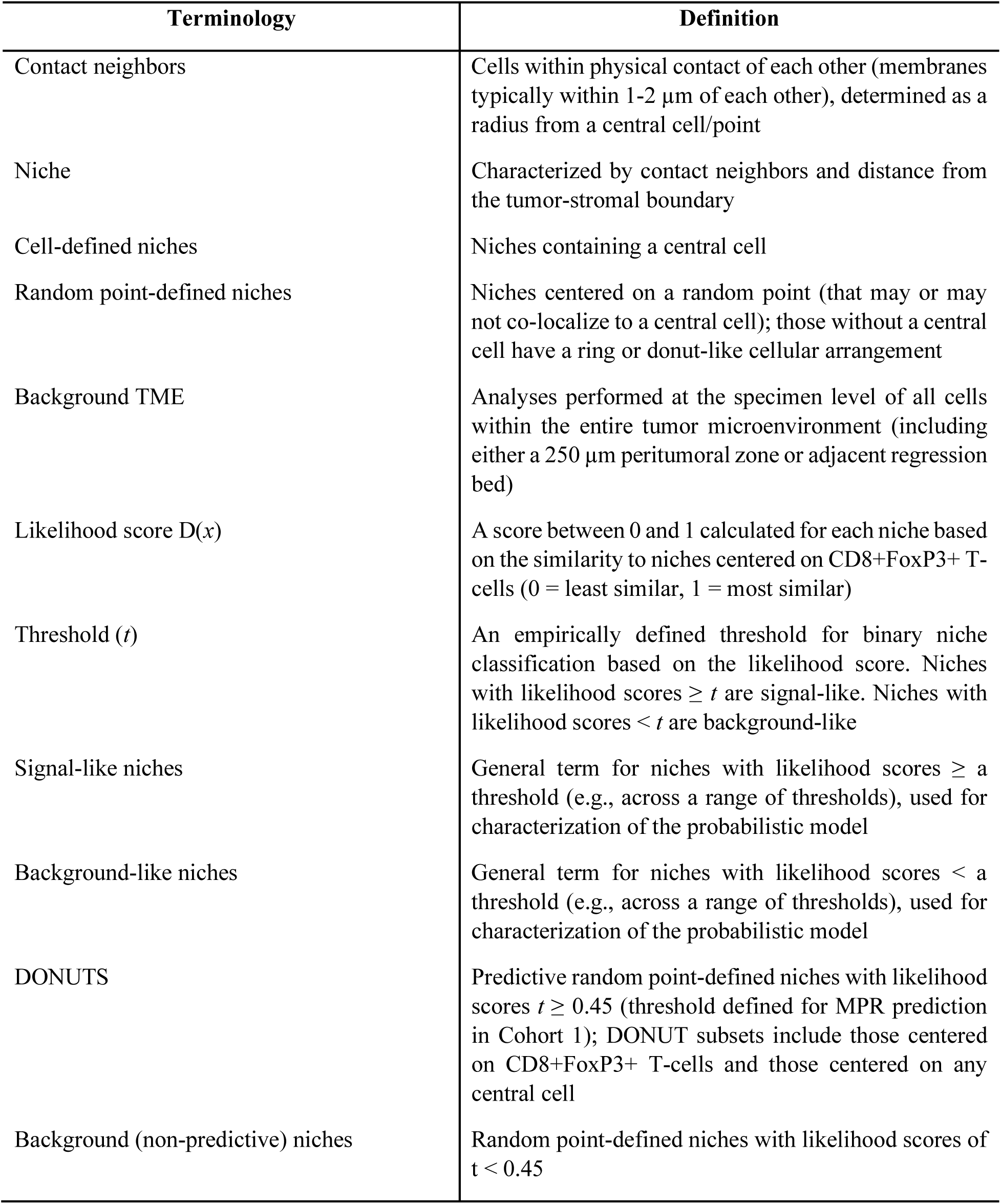
Terminology used for describing spatial biology in the tumor microenvironment.

| Terminology | Definition |
| --- | --- |
| Contact neighbors | Cells within physical contact of each other (membranes typically within 1-2 $\mu\text{m}$ of each other), determined as a radius from a central cell/point |
| Niche | Characterized by contact neighbors and distance from the tumor-stromal boundary |
| Cell-defined niches | Niches containing a central cell |
| Random point-defined niches | Niches centered on a random point (that may or may not co-localize to a central cell); those without a central cell have a ring or donut-like cellular arrangement |
| Background TME | Analyses performed at the specimen level of all cells within the entire tumor microenvironment (including either a 250 $\mu\text{m}$ peritumoral zone or adjacent regression bed) |
| Likelihood score $D(x)$ | A score between 0 and 1 calculated for each niche based on the similarity to niches centered on CD8+FoxP3+ T-cells (0 = least similar, 1 = most similar) |
| Threshold ($t$) | An empirically defined threshold for binary niche classification based on the likelihood score. Niches with likelihood scores $\geq t$ are signal-like. Niches with likelihood scores $< t$ are background-like |
| Signal-like niches | General term for niches with likelihood scores ≥ a threshold (e.g., across a range of thresholds), used for characterization of the probabilistic model |
| Background-like niches | General term for niches with likelihood scores $<$ a threshold (e.g., across a range of thresholds), used for characterization of the probabilistic model |
| DONUTS | Predictive random point-defined niches with likelihood scores $t \geq 0.45$ (threshold defined for MPR prediction in Cohort 1); DONUT subsets include those centered on CD8+FoxP3+ T-cells and those centered on any central cell |
| Background (non-predictive) niches | Random point-defined niches with likelihood scores of $t < 0.45$ |

### Developing a spatial biomarker to predict anti-PD-1 response

We hypothesized that if the CD8+FoxP3+ T-cells were a biomarker of a productive anti-tumor immune response, then their niche may also constitute a biomarker signal associated with response to therapy, **Table 1**. To that end, we developed a probability model, inspired by data analysis techniques used in particle physics investigations to identify the CD8+FoxP3+ cell-defined niche, which includes information on the distance of the niche from the tumor-stromal boundary. The resultant donut-shaped spatial arrangements represented configurations of contact neighbors of CD8+FoxP3+ cells. Once characterized, these donut configurations were then studied without requiring the presence of a central CD8+FoxP3+ cell (**Figure 4C**). The likelihood threshold that defined the niches most predictive of major pathologic response was identified using the Discovery cohort, *i.e.,* pre-treatment specimens from Cohort 1. As this is a probabilistic model, there is not just one resultant configuration, and the niches meeting this threshold are collectively referred to as the **d**iversity **o**f **n**iches **u**nlocking **t**reatment **s**ensitivity (DONUTS). At the selected threshold, the biomarker performance of the DONUTS was improved relative to CD8+FoxP3+ T-cells (AUC=0.79 vs. 0.75 for MPR, respectively, **Figure 4D**). The resultant DONUTS were then visualized using AstroPath, where they were found to concentrate at the tumor-stromal boundary, consistent with an active anti-tumor immune response (**Figure 4E**). Adding further support to the concept that DONUTS represent immunoactive niches, all immune subsets identified by the mIF assay were significantly enriched in the DONUTS compared to non-predictive background niches (**Figure 4F**) and the background TME (**Figure 4G, Figure S5F-J)**.

Although the DONUTS were defined using cell lineage markers only, we hypothesized that there might be an enrichment of PD-1 and PD-L1 expression consistent with adaptive immune resistance.^5,30,31^ We calculated a composite DONUTS configuration that included PD-1 and PD-L1 expression (**Figure 5A, 5B)**. While an ensemble of potential configurations is identified using the probability model, the one shown represents the proportion of evaluated cell lineages and PD-1/L1 expression patterns included in at least 50% of the DONUTS.

**Figure 5.**
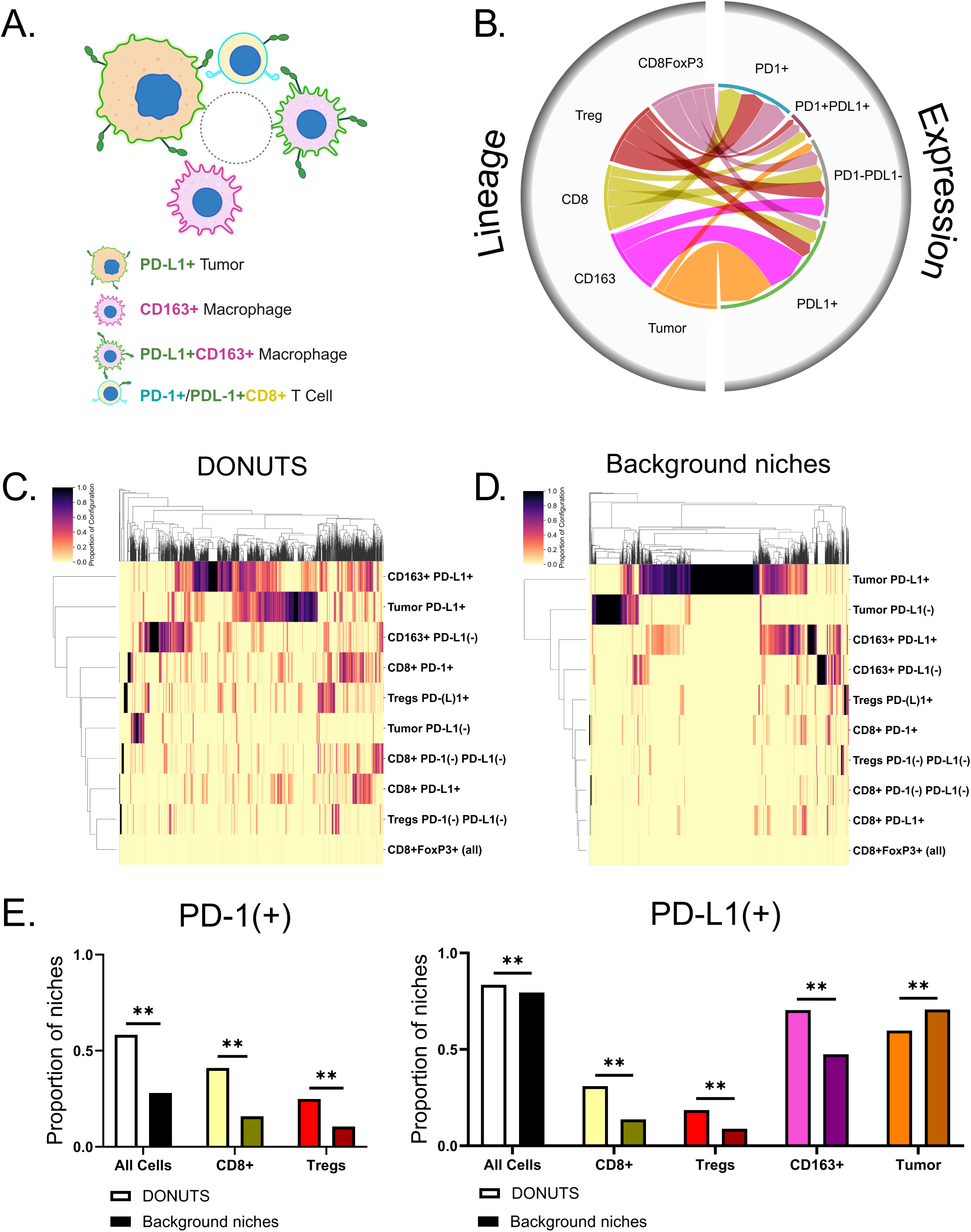
DONUTS are enriched in co-localized cells expressing PD-1 and PD-L1. **(A)** Composite representation of the spatial model showing the cells present in at least 50% of the DONUTS. **(B)** Chord diagram showing the proportional expression of PD-1 and PD-L1 for each cell lineage within the DONUTS. **(C-D)** Heat maps depicting hierarchical clustering of niches by composition, including cellular subsets defined by lineage markers and PD-1 or PD-L1 expression. DONUTS are enriched in PD-L1+/-macrophages, PD-L1+tumor cells, and PD-1+CD8+ T cells. Non-predictive, background TME niches are dominated by tumor cells, including PD-L1+ and (-) subsets. (**E**) The proportion of PD-1+CD8+ T cells and T-regs, as well as PD-L1+CD8+ T cells, T-regs, and CD163+ macrophages are significantly enriched in the DONUTS relative to background TME niches. (**p<0.005)

Hierarchical clustering was used as an orthogonal approach to evaluate PD-1+ and PD-L1+ cellular subsets in DONUTS as compared to background (non-predictive) cellular niches. The percentages of CD8+ T cells expressing PD-1 and PD-L1 were significantly enriched in the DONUTS, as well as the percentage of PD-L1+ CD163+ macrophages (**Figures 5C-E**). In contrast, PD-L1+ tumor cells were not enriched in DONUTS, further underscoring the distinct nature of these two biomarkers. The density of DONUTS positively correlated with the density of PD-1 to PD-L1 proximity metrics (p=0.003), further highlighting spatial, biologic features of the niche underpinning response to PD-1/PD-L1 therapeutic blockade.

### The DONUTS model predicts response to therapy in advanced NSCLC

The association between the densities of DONUTS in pre-treatment biopsies and survival outcomes was then demonstrated in early-stage disease and validated in advanced NSCLC cohorts (**Figure 6A**, **Figure S6**). In Cohort 3, where long-term survival data were not available, the DONUTS density was more strongly associated with radiographic response than the CD8+FoxP3+ T-cell density (AUC 0.86 vs. AUC 0.73, respectively, **Figure S6**). Following neoadjuvant anti-PD-1 therapy, on-treatment DONUT densities correlated positively with TLS density (p=0.003) and negatively with % residual viable tumor as a continuous variable (p=0.001), aligning with known features of pathologic response and underscoring the DONUTS as a representation of an organized, productive anti-tumor immune response.

**Figure 6.**
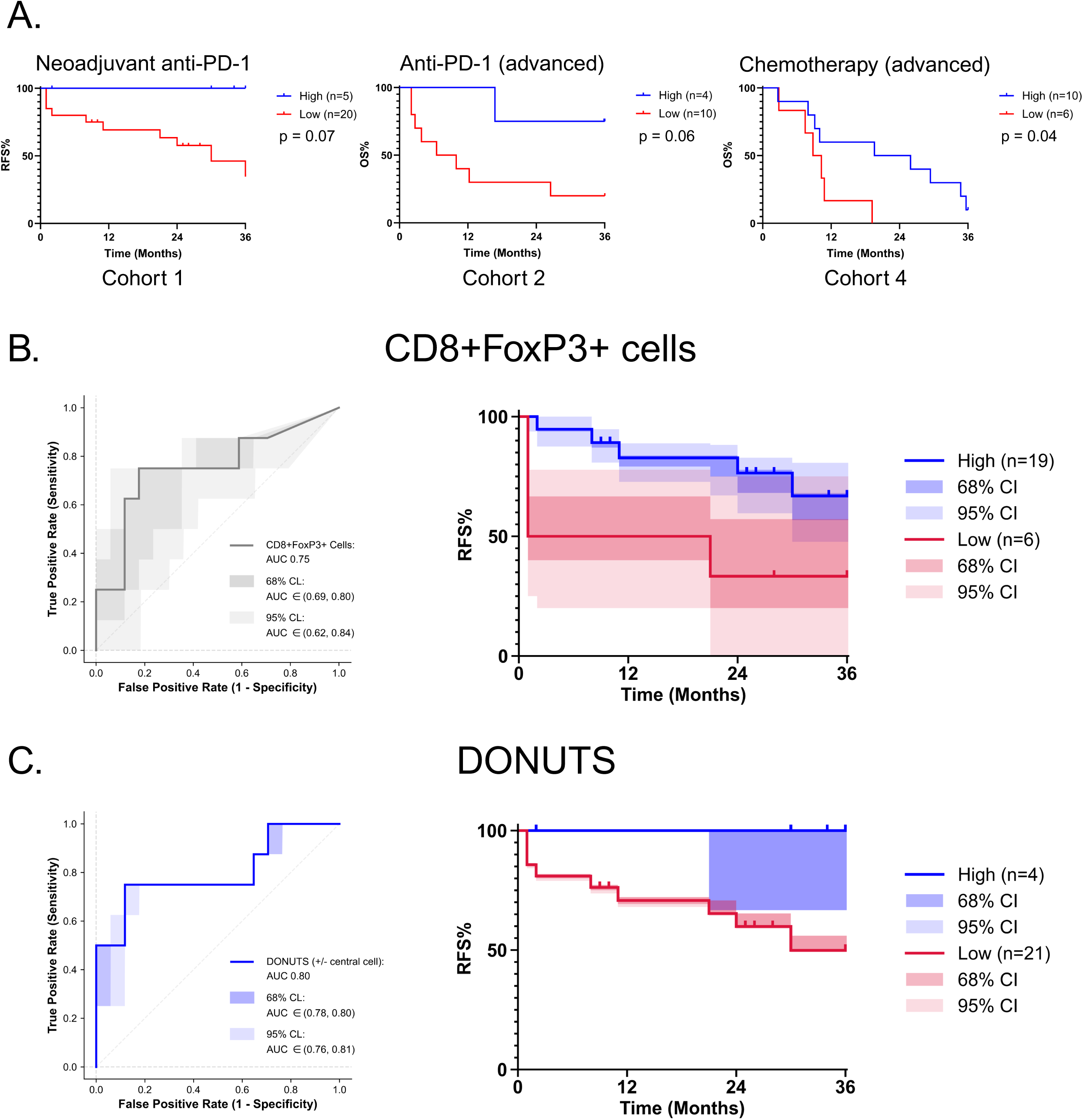
The DONUTS model is a spatial, pre-treatment biomarker that predicts patient outcomes to therapy in both the neoadjuvant and advanced disease settings. **(A)** Pre-treatment DONUTS densities stratify RFS in patients with early-stage NSCLC treated with neoadjuvant anti-PD-1-based therapy (Cohort 1) as well as OS in patients with advanced, pre-treated NSCLC receiving anti-PD-1 therapy (Cohort 2) and chemotherapy (Cohort 4). Also see **Figure S6** for OS for Cohort 1 and PFS for Cohorts 2 and 4. Statistical errors on ROC and KM curves show the impact of analyte abundance biomarker performance of **(B)** CD8+FoxP3+ T-cells themselves vs. **(C)** DONUTS.

To compare the statistical uncertainty associated with predicting treatment outcomes using a conventional vs. spatial biomarker, we next determined the uncertainty on the AUC curves and Kaplan-Meier curves for CD8+FoxP3+ T-cells and the DONUTS. There was a four-fold reduction in statistical uncertainty due to analyte-based uncertainty for predicting MPR in Cohort 1 with the DONUTS relative to CD8+FoxP3+ T-cells (AUC 95% CI [0.62, 0.84] vs. [0.76, 0.81], respectively, **Figure 6B-C**). This difference may be attributed to biomarker prevalence in the TME – the DONUT occurred at 150-fold higher abundance than CD8+FoxP3+ T-cells themselves. The higher abundance of the DONUTS also resulted in a similar reduction in uncertainty on the Kaplan-Meier curves, without a loss of predictive performance.

## DISCUSSION

This study leveraged the substantial technical advances of the AstroPath platform for whole slide mIF to analyze over 35 million cells in the TME of NSCLC. The resultant tumor-immune maps include four patient cohorts that span disease stages and treatment types. We used these maps to identify associations between pre-treatment densities of CD8+FoxP3+ T-cells and their niche with treatment outcomes, including pathologic response and patient survival. To our knowledge, we are also presenting the first whole slide cellular maps showing select immune cell dynamics under therapeutic pressure in the neoadjuvant setting.^22^ Collectively, these spatial data were used to discover and develop a next-generation mIF biomarker (the DONUTS assay) with prognostic significance, which is now positioned for refinement and extended validation in large retrospective datasets, followed by deployment in prospective clinical trials to guide ICB therapy. In addition to initial validation across multiple patient cohorts, the spatial modeling approach described herein reduced statistical uncertainty in performance.

The localization of the CD8+FoxP3+ cells to the tumor-stromal boundary on the tumor-immune maps combined with the robust positive association with patient outcomes led us to hypothesize that this cell phenotype represented an anti-tumor immune cell, rather than a CD8+ variant of a conventional, immunosuppressive CD4+ regulatory T-cell. Profiling of tumor infiltrating CD8+FoxP3+ T-cells by scRNAseq did, in fact, demonstrate a cytotoxic, proinflammatory, and activated phenotype that is also distinct from previously reported double positive CD4+CD8+ T-cell subsets.^32^ Compared to CD4+ Treg populations, CD8+FoxP3+ T-cells demonstrated increased expression of CXCL13, 4-1BB/CD137, and CD39 – all markers of neoantigen-specific CD8+ T-cells.^26,33,34^ While some studies have suggested that FoxP3 expression should be the *sine qua non* of suppressive function,^35^ the few functional studies on CD8+cells displaying FoxP3 in the TME suggest otherwise. For example, one report characterizing tumor infiltrating lymphocytes (TIL) in a GM-CSF secreting vaccine study in mice found that FoxP3 expression is dependent on the local cytokine milieu, and concluded that CD8+FoxP3+ cells are emblematic of a host environment supportive of an effective anti-tumor T-cell response.^36^ There may also be a functional benefit to FoxP3 expression in CD8 cells. In a melanoma model of adoptive T-cell therapy, FoxP3 overexpression in memory-like and effector CD8 T-cells resulted in enhanced metabolic activity enabling cellular proliferation even in glucose limited environments.^37^ While it is possible that bona fide CD8+FoxP3+Tregs exist in other pathologic conditions such as autoimmunity or chronic infection, we find that within the TME, these cells do not have functional programs consistent with immunosuppression. Rather, they represent highly-potent early, effector T-cells, explaining their strong positive association with immunotherapy treatment response.

One of the historic barriers to detecting and studying CD8+FoxP3+ T-cells is that they are relatively rare. CD8+FoxP3+ T-cells represent approximately 0.4% and 0.1% of peripheral blood T-cells in humans and mice, respectively.^20,38^ Within the TME, prevalence rates of 0.2-5.2% of all CD8+ cells have been reported in melanoma, NSCLC, and cervical carcinoma.^20,36,39,40^ Incorporating principles from the data-intensive field of Astronomy,^21^ whole slide imaging using the AstroPath platform (vs. subsampling areas from each slide) produces comprehensive and broad foundational reference maps of the TME, enabling detection of such rare events with statistical certainty.

Across most spatial profiling studies to date, practical constraints have required trade-offs between the number of markers assessed and the size of the tissue areas profiled.^12,41^ Prior studies using higher-plex technologies (∼30-40 markers) have relied on tissue subsampling through tissue microarray (TMA) construction (e.g. 0.6mm diameter tumor core) or region of interest (ROI)/high power field (HPF)-based slide scanning (e.g. 3 fields each 500μm x 500μm in size, totalling 0.75mm^2^ surface area studied per patient).^18,42–44^ Studies using such high-plex assays have found that only 5 or 6 of the markers are necessary for predicting outcomes, with no added benefit to including additional markers.^18^ One high-plex study using a TMA format required the use of 12 markers (out of 43 tested) and *both* pre- and on-treatment specimens to demonstrate spatial determinants associated with a response to immunotherapy, achieving an AUC of 0.8 in a discovery cohort (this study did not include a validation cohort).^17^ In contrast, we are able to demonstrate a predictive AUC of 0.8 that validates in multiple cohorts using 4 markers by mapping more of the TME. Specifically, the breadth of the TME covered is on average 15x greater in pre-treatment specimens and ∼100x greater for on-treatment specimens. Given that the DONUTS biomarker is additive to PD-L1 IHC, PD-L1 could also readily be included as a fifth marker in the pre-treatment mIF assay.

The use of a reduced number of markers and characterization of the whole slide image are important features that lend themselves to clinical implementation. Importantly, while some of the higher plex methodologies are useful for efficiently examining more markers for research purposes, they are also not poised for clinical deployment, and TMA and HPF/ROI-based tissue input strategies are also not clinical fodder. Chromogenic IHC is the gold standard in pathology for clinical diagnostics, and the mIF 4-to 6-plex assays reported herein have been rigorously optimized and undergone quantitative validation against chromogenic IHC according to best-practice guidelines.^12^ Additionally, whole slide mIF has tissue inputs and staining platforms that mirror current IHC workflows. Further, a version of the 6-plex mIF assay described herein has also been shown to be highly reproducible in a multi-institutional study.^45^

While whole slide images of pre-treatment tissue biopsies are meaningfully larger than the TMA spots and select HPF sampling used in other spatial studies, sampling error related to the detection of a rare cell type could still be a concern for predictive diagnostic assays. The data-rich TME maps were used to enable identification of distinct cellular niches surrounding CD8+FoxP3+ T-cells that were detected across multiple patient cohorts and stages of NSCLC. Our DONUTS model effectively represents these niches even in the absence of a central CD8+FoxP3+ cell, and we found that the DONUTS biomarker was 150-fold more abundant than CD8+FoxP3+ T-cells themselves, enhancing sensitivity and reducing sampling error in small specimens.

It is also worth emphasizing the association of CD8+FoxP3+ cells and DONUTS densities with the number of TLS. While TLS have been associated with immunotherapy response, they have been characterized and studied in on-treatment specimens.^8,9,46^ They are also not often sampled in pre-treatment lung biopsies, as they tend to be located at the perimeter of the TME.^9^ Here, we demonstrate an association between the CD8+FoxP3+ cells/DONUTS that can be visualized in pre-treatment specimens and the presence of TLS, thus bringing a TLS-related metric into a predictive setting.

In addition to defining a predictive immunoactive niche for ICB therapy, we unexpectedly identified associations between CD8+FoxP3+ T-cells and their niche with OS and PFS for patients receiving second-line chemotherapy (docetaxel) for advanced NSCLC. It is recognized that significant overlap exists between predictive and prognostic signals for immunotherapy biomarkers (*e.g.*, CD8 density and/or PD-L1 expression).^30,47–49^, Although we cannot entirely exclude the possibility that CD8+FoxP3+ T-cells and their niches may be prognostic biomarkers, a prior study of N=87 patients with NSCLC across multiple stages found that CD8+FoxP3+/CD8+ T-cell frequency is not associated with survival.^39^ Rather, emerging evidence suggests an immunostimulatory role for some traditional chemotherapies, and these patients had received docotaxel in the first line, which may have immunologically primed the TME.^50^ Historically it was thought that chemotherapy would create an immunosuppressive environment that would negatively impact immunotherapy. Instead, animal studies demonstrate docetaxel-mediated TME remodeling that includes enhanced antigen-specific T-cell responses^51,52^. Further, a study of docetaxel-based chemohormonal therapy in patients with prostate cancer confirmed that relative to pre-treatment biopsies, on-treatment biopsies showed increased T-cell infiltration and upregulation of PD-1 and PD-L1.^51^ The immunologic biomarker signals identified in the current study add to provocative evidence for an immunomodulatory role for select traditional chemotherapies that may help explain the synergistic efficacy of some immunochemotherapy regimens in patients with NSCLC.^1–3^

Limitations of this study include the small cohort sizes, though the fact that the biomarker signal validated in multiple cohorts is encouraging. Most of the cohorts included in this study were derived from practice-changing clinical trials, and remaining tissue available for study was only available for a subset of patients. Additionally, while we included patients with early- and late-stage NSCLC treated with anti-PD-1 or chemotherapy, we did not study patients treated with chemoimmunotherapy with mIF, though it is anticipated that our findings would also apply to that treatment setting. We did perform scRNAseq from patients treated with chemoimmunotherapy, and showed that CD8+FoxP3+ cells identified in these patients demonstrated the same signature as seen in patients treated with ICB alone. Further, a recent single cohort study described the association of pre-treatment CD8+FoxP3+ cells with patient outcomes small group of patients with advanced NSCLC treated with chemoimmunotherapy, suggesting the applicability of the DONUTS biomarker in this setting.^53^ Notwithstanding, one setting where this assay could potentially be used would be to select for those patients who would benefit from anti-PD-1 alone, avoiding chemotherapy.^54^ Another potential application could be to help identify responders to PD-1-based therapies in the PD-L1(-) patient subgroup.^53^ Future studies of large patient cohorts are required for extended analytic and clinical validation of specific assay conditions (e.g., to determine the optimal thresholds for patient stratification, which will likely vary by squamous versus non-squamous tumors and by treatment setting).

In summary, the DONUTS model is a computational, spatial biomarker that reflects the unique cellular niche of CD8+FoxP3+ cells, which we demonstrate represent highly activated, effector T-cells. DONUTS represents an advance beyond other early candidate, spatial biomarkers ^17–19,44,55^ in that it is performed on whole slides, uses a mIF assay that has been optimized to clinical standards, requires a limited number of markers, is performed on pre-treatment specimens only, predicts response to therapy, and validates as biomarker signal across multiple cohorts. CD8+FoxP3+ T-cells have now been identified in the pre-treatment TME of patients responding to anti-PD-1 based therapies across multiple tumor types and stages,^20,56^ raising consideration for their utility and/or the utility of DONUTS as a potential pan-tumor, pre-treatment biomarker. This computational model could not have been developed without the use of highly accurate tumor-immune maps with single-cell resolution across the wider TME. To facilitate usage of these data for the greater research community, we present these resultant Atlases and associated clinical data as a new science domain on SciServer (https://www.sciserver.org/integration/astropath/).^57^ The web address links to SciServer, a Big Data analytics platform containing several spatial databases for astronomy, turbulence, ocean circulation, and now with a new chapter for the tumor microenvironment.

## RESOURCE AVAILABILITY

Cell density and DONUT density tables on an individual patient basis can be found at https://data.mendeley.com/preview/4b6cwdsp7p?a=379b61e4-757b-48b1-a0fa-3e8155c84956

(DOI: 10.17632/4b6cwdsp7p.2; embargoed until publication). Computer code for the DONUTS algorithm can be found at https://github.com/AstroPathJHU/DONUT.

## ACKNOWLEDGEMENTS

The authors would like to acknowledge Jose Loyola and Bhakti Pandey for excellent technical assistance, as well as Dr. Robin Edwards and Darren Locke (both previously at Bristol Myers Squibb) and Cliff Hoyt (Akoya/Quanterix). This work was supported by The Mark Foundation for Cancer Research (JMT, AS, KNS, DMP); Sidney Kimmel Cancer Center Core Grant P30 CA006973 (JMT, LD); National Cancer Institute R01 CA142779 (JMT, DMP); NIH T32 CA193145 (JSD); R50 CA243627 (LD); NIH R37 CA251447 (KNS); NIH R01 HG013409 (HJ); and The Bloomberg-Kimmel Institute for Cancer Immunotherapy. Patients treated at Memorial Sloan Kettering Cancer Center were supported in part by Memorial Sloan Kettering Cancer Center Support Grant/Core Grant (P30 CA008748). This study was conducted with the support of the Ontario Institute for Cancer Research through funding provided by the Government of Ontario (TRC). This research makes use of the SciServer science platform (www.sciserver.org). SciServer is a collaborative research environment for large-scale data-driven science. It is being developed at, and administered by, the Institute for Data Intensive Engineering and Science at Johns Hopkins University. SciServer is funded by the National Science Foundation through the Data Infrastructure Building Blocks (DIBBs) program and others, as well as by the Alfred P. Sloan Foundation and the Gordon and Betty Moore Foundation.

## AUTHOR CONTRIBUTIONS

Conceptualization: T.R.C, J.S.R., B.F.G., A.S.S., J.M.T., J.C.S.;

Data curation: T.R.C., J.M.T., J.R.S., B.F.G., M.F., E.C.;

Formal analysis: J.S.R., M.F., E.C., T.R.C., J.M.T.;

Funding acquisition: J.M.T., A.S.S., D.M.P., K.N.S.;

Investigation: T.R.C., J.M.T., J.S.R., M.F., E.C., D.W., S.U., K.P., D.V., M.M., A.F., N.E., T.P., B.Z., Z.Z., J.X.C., J.Z., J.S., L.L.E., N.D.;

Methodology: T.R.C., J.M.T., A.S.S, J.R.S., M.F., B.F.G., E.W., J.S.D.;

Project administration: T.R.C., J.M.T., M.E., H.J.;

Resources: T.R.C., J.M.T., A.S.S., J.E.C., J.R.B., M.C., J.E.R., P.M.F.;

Software: A.S.S., J.S.R., S.S.D., M.E., S.T., A.J., A.W., D.J.S., D.M., B.F.G., M.F.;

Supervision: J.M.T., A.S.S., T.R.C., D.M.P., K.N.S.;

Validation: J.R.S., M.F., T.R.C., J.M.T., L.L.E., A.O.;

Visualization: M.F., J.R.S., T.R.C., J.M.T.;

Writing – original draft: T.R.C., J.M.T., J.R.S., B.F.G., M.F.;

Writing – review and editing: *All authors.

## METHODS

No statistical methods were used to predetermine sample size. The experiments were not randomized. The investigators were not blinded to allocation during experiments and outcome assessment.

### Patient specimens and clinical course

This study was approved by the Institutional Review Boards (IRB) at Johns Hopkins University (JHU) and Memorial Sloan Kettering Cancer Center and was conducted in accordance with the Declaration of Helsinki and the International Conference on Harmonization Good Clinical Practice guidelines. Written informed consent was obtained from all patients at the time of study enrollment.

Cohort 1 was derived from the first-in-human clinical trial of neoadjuvant anti-PD-1-based therapy in patients with resectable non-small cell lung cancer (NCT02259621).^22,58^ Newly diagnosed patients with resectable, stage IA – IV NSCLC were treated with anti-PD-1 based therapy prior to definitive surgical resection. Patients received two to three doses (3mg/kg) of either nivolumab or pembrolizumab alone (n=19) or in combination with one dose (1mg/kg) of ipilimumab (n=6). Therapy was administered over 8 weeks followed by surgical resection performed within 10 days of the last dose. Demographic and clinical data were collected (**Table S1**). Recurrence-free survival was calculated using the date of enrollment to the date of clinical recurrence or last follow-up in patients who did not recur.

Archival formalin-fixed paraffin-embedded (FFPE) pre-treatment tumor tissues (n=42) from n=29 unique patients were obtained from the surgical pathology archives at Johns Hopkins Hospital and Memorial Sloan Kettering Cancer Center. The final cohort consisted of n=25 specimens that met inclusion criteria from n=25 unique patients (**Figure S1A**). Pathologic response in the corresponding on-treatment resection specimens was previously reported.^9,22,58^ Pathologic response was determined by scoring the percent residual viable tumor (%RVT) at 10% intervals.^9,59,60^ Patients whose on-treatment tumors contained ≤10% RVT were classified as major pathologic responders (MPR), and those with ≤50%RVT as partial pathologic responders. On-treatment FFPE tumor resection specimens (n=35) were also evaluated (**Figure S1B**), including n=23 paired pre- and on-treatment specimens from the same patient. In a subset of patients, fresh tumor tissue was collected at the time of surgical resection for single-cell RNA sequencing (n=15).

Cohort 2 consisted of patients with advanced, pre-treated non-squamous NSCLC who received anti-PD-1 therapy as part the CheckMate 057 clinical trial (NCT01673867).^23^ Of the n=21 pre-treatment tumor tissues available, n=14 met inclusion criteria for Cohort two (**Figure S1C**).

Cohort 3 included patients with advanced NSCLC who progressed on first-line chemotherapy and received standard of care second-line anti-PD-1 therapy). All patients with pre-treatment tumor tissue available (n=20) met inclusion criteria (**Figure S1D**).

Cohort 4 consisted of patients with advanced, pre-treated non-squamous NSCLC who received second-line docetaxel chemotherapy in the control arm of CheckMate 057 (NCT01673867) ^23^. Of the n=22 pre-treatment tumor tissues available, n=16 met inclusion criteria for Cohort four (**Figure S1E**).

### Multiplex immunofluorescence staining

For Cohort one, a six-plex mIF panel (PD-1, PD-L1, CD8, FoxP3, CD163, pan-cytokeratin) was quantitatively validated against chromogenic IHC in NSCLC specimens (**Figure S2**). Briefly, four-micron thick FFPE slides underwent sequential staining cycles. Primary and secondary antibodies are described in **Table S5.** Signals were visualized with Opal fluorophores (Akoya Biosciences). To normalize staining intensity across batches, each staining batch included a tissue microarray (TMA) control slide.^20^ For the advanced NSCLC cohorts (cohorts 2-4), versions of this assay where used that substituted CD68 for CD163 as a macrophage marker.

### Image acquisition, processing, and analysis

A Vectra Imaging System (Akoya Biosciences, MA) was used to acquire whole slide survey scans at 10x (1.0µm/pixel) magnification (.QPTIF file). Twenty percent overlapping high powered field (HPF) image tiles at 20x (0.5µm/pixel) magnification were acquired for the entire tissue area using multispectral scanning (.im3 files). Annotations were performed on the survey scans using HALO (Indica Labs, NM) by a board-certified pathologist to delineate the tumor boundary and exclude necrosis and staining artifacts. The peri-tumor region was defined as the 250 μm into the stroma beyond the tumor-stromal boundary. For the on-treatment specimens, the tumor bed was annotated to include both the residual tumor and the regression bed, i.e. the original tumor extent prior to regression.^9,59,61^

Cell segmentation maps and single-cell image analysis data were generated using a multi-pass deployment of inForm v2.4.8 (Akoya Biosciences), and visually verified as previously described.^20^ Individual HPF tiles and associated single-cell data were assembled into a seamless whole slide image with an absolute coordinate system at sub-micron resolution.

The fully processed mIF images, annotations, and cellular data were loaded into the AstroPath relational (SQL) database with associated clinical and pathology data, which was formatted using AstroID, a 6-tier hierarchical data structure.^62^ Instructions for accessing the images in the AstroPath database are described in **Table S4**. For all pre-treatment specimens densities for all cell lineages were calculated for cells within the TME, defined as the intra-tumoral region plus the peritumoral zone. One pre-treatment specimen was a lymph node metastasis and only the intra-tumoral area was analyzed. For on-treatment specimens, densities were calculated over the tumor bed or the intra-tumoral region plus the peritumoral zone if no regression was present. High vs. low density thresholds were defined for each cohort using the Youden’s index. PD-1 to PD-L1 proximity was calculated by determining the density of PD-L1+ cells within 25µm of a PD-1+ cell.^63^

### Identification and biomarker value of immediate topology surrounding CD8+FoxP3+ T-cells: development of DONUTS

We designed a method to assign a likelihood score that quantifies how closely a given cell’s spatial neighborhood resembles that of a CD8+FoxP3+ cell, accounting for its relative distance to the tumor-stromal boundary. Definitions of terminology used to describe spatial arrangements is presented in **Table 1**. This approach draws inspiration from methods in particle physics, where likelihood scores are used as *discriminants* to distinguish between two competing hypotheses, *A* and *B*.^64^ There the likelihood score, *d(x)*, is derived from the Neyman-Pearson lemma, which states that, given an observation *x* and two hypotheses *A* and *B*, the most powerful test for distinguishing between them is the ratio of their likelihoods.^65^ That is:

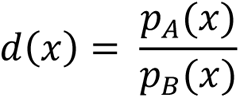

A commonly used equivalent formulation expresses the likelihood as a normalized score, *D(x)*, called *discriminant* in particle physics:

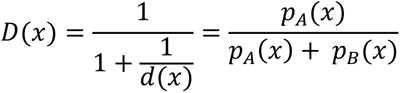

This formulation is convenient because it is bounded within the interval [0, 1], where *D(x)∼1* for an observation more likely to have been produced by hypothesis *A* and *D(x)∼0* for an observation more likely to have been produced by hypothesis *B*. In this case, we take hypothesis *A* to be that the cell’s spatial neighborhood resembles the spatial neighborhood of a CD8+FoxP3+ cell at a similar relative distance to the tumor-stromal boundary and hypothesis *B* is that it does not (i.e., it resembles some other cell type’s neighborhood at that relative distance). Thus, the likelihood score quantifies how similar the arrangement is to one that typically surrounds a central CD8+FoxP3+ cell, with larger values meaning that the neighborhood looks more like a CD8+FoxP3+ neighborhood. Furthermore, the likelihood score reflects the degree of niche similarity regardless of the actual cell type of the central cell or whether there is even a central cell at all, while also accounting for the cell’s relative distance to the tumor-stromal boundary.

We define each cell’s niche (*x*) by measuring (1) the cell’s location relative to the tumor-stromal boundary and (2) the lineage distribution of each cell’s contact neighbors. The distance from the tumor-stromal boundary was assessed in 25µm bins, with negative values for intratumoral cells and positive values for peritumoral cells. Cellular contact neighbors were denoted as the cells within 5 – 10µm around each cell. The lower limit of 5μm was chosen so that the cell at the center of the arrangement was not itself counted. The number of contact neighbors for each cell lineage was similar across all lineages evaluated (**Figure S7A**).

To evaluate for *D*(*x*), we established empirical distributions for the features of CD8+FoxP3+ cell niches [*p*_A_(*x*)] and the niches of all other cell lineages [*p*_B_(*x*)] using six on-treatment NSCLC tissue samples stained with the same mIF panel. These specimens were selected because each contained over 1,000 CD8+FoxP3+ T-cells in the analysis area (tumor plus 250 µm adjacent stroma). This training data (12.8M total cells and ∼50,000 CD8+FoxP3+ T-cells) was used to define features that distinguish CD8+FoxP3+ T-cell niches from those of all other cell lineages. We estimated p_A (x) as the fraction of CD8+FoxP3+ cells in the training set that have niche x and p_B (x) as the fraction of cells of other lineages that have niche x. Subsequently, the likelihood score [*D*(*x*)] for each cell in the test datasets was calculated by comparing the features of the cell’s niche (*x*) to the empirical distributions defined with the training data.

After *D(x)* was defined for each cell, we applied a range of thresholds (*t*) to classify niches in the pre-treatment specimens from Cohort 1 as either more similar to the CD8+FoxP3+ T-cell niche (referred to as signal-like niches, *D(x)* ≥ *t*) or background-like niches (*D(x)* < *t*). We then examined how the distribution of central cell lineages within predicted signal-like niches varied across threshold values. As the threshold increased, the proportion of signal-like niches with a CD8+FoxP3+ central cell also increased relative to those centered around other cell lineages (**Figure S7B**).

To characterize niches independent of the presence of a central cell, we sampled random locations to use as the centroid to evaluate for *D(x)* (i.e., random point-defined-niches). These points were generated at an average density of 8,000 points per HPF (∼5,900 points per mm^2^), matching the average cell density observed in the TME. This allows for a direct comparison between the densities of random point-defined niches and cell-defined niches. We evaluated *D(x)* at each random-point and measured the proportion of signal-like niches (*D(x)* ≥ t) across the same range of thresholds in Cohort 1. As the threshold increased, the proportion of signal-like niches centered on random points declined (**Figure S7B**). This mirrored the trend observed for niches centered around non-CD8+FoxP3+ cell lineages, demonstrating selective enrichment of niches centered on CD8+FoxP3+ cells as the likelihood score increases.

To quantify the ability of signal-like niche density to distinguish responders from non-responders across varying thresholds, we used the random point defined-niches to generate receiver operating characteristic (ROC) curves and associated area under the curve (AUC) estimates for patients in Cohort 1. We found that using a threshold of *t ≥ 0.45* provided both strong predictive performance for treatment response (**Figure S7C**) and a sufficient density of signal-like niches to ensure reliable detection (**Table S7**). In all subsequent analyses (including those of Cohorts 2, 3, and 4), the “DONUTS” biomarker is assessed as the density of signal-like niches (centered on random point spatial locations) that have a discriminant scores (*D(x)*) at or above this threshold (*t ≥ 0.45*). Cellular configurations (centered on random point spatial locations) with discriminant scores (*D(x)*) below this defined threshold (*t<0.45*) are referred to as “Background niches”. High vs. low DONUTS density thresholds were defined for each cohort using the Youden’s index

### Establishing analyte-based uncertainty

Monte-Carlo simulations were performed to estimate the analyte-based uncertainty in predicting patient outcomes in Cohort 1. The densities of CD8+FoxP3+ T-cells and DONUTS were simulated for each sample as Poisson distributions. For instance, if a sample contained 100 CD8+FoxP3+ cells distributed over a 10 mm^2^ tissue area, we modeled the number of cells using a Poisson distribution with λ = 100, corresponding to an expected density of 10 cells per mm^2^. This approach yields a two-tailed density distribution, where approximately 70% of simulations fall between 9 and 11 cells per mm^2^. We performed 10,000 Monte-Carlo trials, independently sampling from the Poisson distribution for each trial to generate 10,000 simulated ROC curves. From these, we computed the 68% and 95% confidence intervals for the corresponding ROC using empirical quantiles (**Figure 6B-C**).

For the Kaplan-Meier curves, we used a likelihood-based approach derived from methods used in particle physics. We computed the Poisson negative log-likelihood (NLL) to select patients for the DONUTS high-vs. low-density curves. For each possible Kaplan-Meier probability, all combinations of included and excluded patients that could produce that probability were considered. Gurobi optimization^66^ was used to find the minimum total NLL among all of those combinations. Probabilities that could not be reached with an NLL of less than 1.0 were excluded at 68% confidence level, and probabilities that could not be reached with an NLL of less than 3.84 were excluded at 95% confidence level.

### Single-Cell RNA Sequencing

Gene expression profiles were obtained from scRNAseq datasets presented in Caushi et. al^26^ and Liu et. al^29^, representing specimens from patients treated with anti-PD-1 and anti-PD-1+chemotherapy, respectively. Tumor infiltrating cells with the top five percentile of FoxP3 expression were retained for downstream analysis. Low quality cells with less than 250 genes and genes that were expressed in less than five cells were further filtered. Seurat v4^67–70^ was used to normalize the raw count data, identify top 2000 highly variable genes (HVGs), scale features, and perform principal component analysis (PCA). Harmony algorithm^71^ was applied to integrate across samples and batches. Immunoglobulin genes, gene features associated with type I Interferon response and specific mitochondrial related genes were excluded from clustering. Cell markers were identified using two-sided Wilcoxon rank sum test with adjusted p-value ≤ 0.05. Clusters were labeled based on top differential genes and canonical immune cell markers.

To further interrogate the CD8+FoxP3+ cell subpopulation from Caushi et. al ^26^, cells from cluster 5 were extracted (n=1,299 cells, with high CD8 expression, **Figures 3A-B**) and refined clustering was performed (**Figure 3C**). The processing pipeline was applied as described above (*e.g.*, normalization, HVGs identification and PCA) and Seurat v4 was used to integrate across samples and batches. Cell barcodes from the CD8+ stem-like memory cell subpopulation were obtained from a prior analysis of these samples^26^. A comparison of expression levels between subcluster 0 in **Figure 3C** and the stem-like memory subpopulation was performed by utilizing a two-sided Wilcoxon rank sum test with adjusted p-value ≤ 0.05.

### Statistics

Statistics were performed using Python 3.11^72^ and visualized with Prism (v9.1.0, GraphPad Software, MA). A Shapiro-Wilk test was used to assess the normality of continuous variables. Continuous variables were tested for association with MPR status using one-sided Wilcoxon rank-sum tests, while signed-rank tests were performed to compare densities between paired samples.^73^ Rank-sum values were converted into AUC values.^74^ Associations of immune cell densities with survival were evaluated using Kaplan-Meier curves and log-rank tests. Survival probabilities between groups at the two-year timepoint were evaluated with z-tests. Probabilistic density was calculated by normalizing cell densities at discrete increments from tumor to the average density across all samples at the same distances for a given cell type. Spearman rank correlations were used to evaluate monotonic relationships between continuous variables. P-values less than 0.05 are considered significant.

## REFERENCES

1. Forde, P.M., Spicer, J., Lu, S., Provencio, M., Mitsudomi, T., Awad, M.M., Felip, E., Broderick, S.R., Brahmer, J.R., Swanson, S.J., et al. (2022). Neoadjuvant Nivolumab plus Chemotherapy in Resectable Lung Cancer. N Engl J Med 386, 1973–1985. 10.1056/NEJMoa2202170.

2. Cascone, T., Awad, M.M., Spicer, J.D., He, J., Lu, S., Sepesi, B., Tanaka, F., Taube, J.M., Cornelissen, R., Havel, L., et al. (2024). Perioperative Nivolumab in Resectable Lung Cancer. N Engl J Med 390, 1756–1769. 10.1056/NEJMoa2311926.

3. Wakelee, H., Liberman, M., Kato, T., Tsuboi, M., Lee, S.H., Gao, S., Chen, K.N., Dooms, C., Majem, M., Eigendorff, E., et al. (2023). Perioperative Pembrolizumab for Early-Stage Non-Small-Cell Lung Cancer. N Engl J Med 389, 491–503. 10.1056/NEJMoa2302983.

4. Heymach, J.V., Harpole, D., Mitsudomi, T., Taube, J.M., Galffy, G., Hochmair, M., Winder, T., Zukov, R., Garbaos, G., Gao, S., et al. (2023). Perioperative Durvalumab for Resectable Non-Small-Cell Lung Cancer. N Engl J Med 389, 1672–1684. 10.1056/NEJMoa2304875.

5. Pardoll, D.M. (2012). The blockade of immune checkpoints in cancer immunotherapy. Nat Rev Cancer 12, 252–264. 10.1038/nrc3239.

6. Taube, J.M., Galon, J., Sholl, L.M., Rodig, S.J., Cottrell, T.R., Giraldo, N.A., Baras, A.S., Patel, S.S., Anders, R.A., Rimm, D.L., and Cimino-Mathews, A. (2018). Implications of the tumor immune microenvironment for staging and therapeutics. Mod Pathol 31, 214–234. 10.1038/modpathol.2017.156.

7. Schumacher, T.N., and Thommen, D.S. (2022). Tertiary lymphoid structures in cancer. Science 375, eabf9419. 10.1126/science.abf9419.

8. Fridman, W.H., Meylan, M., Petitprez, F., Sun, C.M., Italiano, A., and Sautès-Fridman, C. (2022). B cells and tertiary lymphoid structures as determinants of tumour immune contexture and clinical outcome. Nat Rev Clin Oncol 19, 441–457. 10.1038/s41571-022-00619-z.

9. Cottrell, T.R., Thompson, E.D., Forde, P.M., Stein, J.E., Duffield, A.S., Anagnostou, V., Rekhtman, N., Anders, R.A., Cuda, J.D., Illei, P.B., et al. (2018). Pathologic Features of Response to Neoadjuvant Anti-PD-1 in Resected Non-Small Cell Lung Carcinoma: A Proposal for Quantitative Immune-Related Pathologic Response Criteria (irPRC). Ann Oncol. 10.1093/annonc/mdy218.

10. Stein, J.E., Soni, A., Danilova, L., Cottrell, T.R., Gajewski, T.F., Hodi, F.S., Bhatia, S., Urba, W.J., Sharfman, W.H., Wind-Rotolo, M., et al. (2019). Major pathologic response on biopsy (MPRbx) in patients with advanced melanoma treated with anti-PD-1: evidence for an early, on-therapy biomarker of response. Ann Oncol 30, 589–596. 10.1093/annonc/mdz019.

11. Lu, S., Stein, J.E., Rimm, D.L., Wang, D.W., Bell, J.M., Johnson, D.B., Sosman, J.A., Schalper, K.A., Anders, R.A., Wang, H., et al. (2019). Comparison of Biomarker Modalities for Predicting Response to PD-1/PD-L1 Checkpoint Blockade: A Systematic Review and Meta-analysis. JAMA Oncol. 10.1001/jamaoncol.2019.1549.

12. Taube, J.M., Akturk, G., Angelo, M., Engle, E.L., Gnjatic, S., Greenbaum, S., Greenwald, N.F., Hedvat, C.V., Hollmann, T.J., Juco, J., et al. (2020). The Society for Immunotherapy of Cancer statement on best practices for multiplex immunohistochemistry (IHC) and immunofluorescence (IF) staining and validation. J Immunother Cancer 8. 10.1136/jitc-2019-000155.

13. Patel, S.S., Weirather, J.L., Lipschitz, M., Lako, A., Chen, P.H., Griffin, G.K., Armand, P., Shipp, M.A., and Rodig, S.J. (2019). The microenvironmental niche in classic Hodgkin lymphoma is enriched for CTLA-4-positive T cells that are PD-1-negative. Blood 134, 2059–2069. 10.1182/blood.2019002206.

14. Magen, A., Hamon, P., Fiaschi, N., Soong, B.Y., Park, M.D., Mattiuz, R., Humblin, E., Troncoso, L., D’souza, D., Dawson, T., et al. (2023). Intratumoral dendritic cell-CD4+ T helper cell niches enable CD8+ T cell differentiation following PD-1 blockade in hepatocellular carcinoma. Nat Med 29, 1389–1399. 10.1038/s41591-023-02345-0.

15. Lopez Janeiro, A., Miraval Wong, E., Jiménez-Sánchez, D., Ortiz de Solorzano, C., Lozano, M.D., Teijeira, A., Schalper, K.A., Melero, I., and De Andrea, C.E. (2024). Spatially resolved tissue imaging to analyze the tumor immune microenvironment: beyond cell-type densities. J Immunother Cancer 12. 10.1136/jitc-2023-008589.

16. Taube, J.M., Sunshine, J.C., Angelo, M., Akturk, G., Eminizer, M., Engle, L.L., Ferreira, C.S., Gnjatic, S., Green, B., Greenbaum, S., et al. (2025). Society for Immunotherapy of Cancer: updates and best practices for multiplex immunohistochemistry (IHC) and immunofluorescence (IF) image analysis and data sharing. J Immunother Cancer 13. 10.1136/jitc-2024-008875.

17. Wang, X.Q., Danenberg, E., Huang, C.S., Egle, D., Callari, M., Bermejo, B., Dugo, M., Zamagni, C., Thill, M., Anton, A., et al. (2023). Spatial predictors of immunotherapy response in triple-negative breast cancer. Nature 621, 868–876. 10.1038/s41586-023-06498-3.

18. Sorin, M., Rezanejad, M., Karimi, E., Fiset, B., Desharnais, L., Perus, L.J.M., Milette, S., Yu, M.W., Maritan, S.M., Doré, S., et al. (2023). Single-cell spatial landscapes of the lung tumour immune microenvironment. Nature 614, 548–554. 10.1038/s41586-022-05672-3.

19. Schürch, C.M., Bhate, S.S., Barlow, G.L., Phillips, D.J., Noti, L., Zlobec, I., Chu, P., Black, S., Demeter, J., McIlwain, D.R., et al. (2020). Coordinated Cellular Neighborhoods Orchestrate Antitumoral Immunity at the Colorectal Cancer Invasive Front. Cell 183, 838. 10.1016/j.cell.2020.10.021.

20. Berry, S., Giraldo, N.A., Green, B.F., Cottrell, T.R., Stein, J.E., Engle, E.L., Xu, H., Ogurtsova, A., Roberts, C., Wang, D., et al. (2021). Analysis of multispectral imaging with the AstroPath platform informs efficacy of PD-1 blockade. Science 372. 10.1126/science.aba2609.

21. Szalay, A.S., and Taube, J.M. (2022). Data-Rich Spatial Profiling of Cancer Tissue: Astronomy Informs Pathology. Clin Cancer Res 28, 3417–3424. 10.1158/1078-0432.CCR-19-3748.

22. Forde, P.M., Chaft, J.E., Smith, K.N., Anagnostou, V., Cottrell, T.R., Hellmann, M.D., Zahurak, M., Yang, S.C., Jones, D.R., Broderick, S., et al. (2018). Neoadjuvant PD-1 Blockade in Resectable Lung Cancer. N Engl J Med 378, 1976–1986. 10.1056/NEJMoa1716078.

23. Borghaei, H., Paz-Ares, L., Horn, L., Spigel, D.R., Steins, M., Ready, N.E., Chow, L.Q., Vokes, E.E., Felip, E., Holgado, E., et al. (2015). Nivolumab versus Docetaxel in Advanced Nonsquamous Non-Small-Cell Lung Cancer. N Engl J Med 373, 1627–1639. 10.1056/NEJMoa1507643.

24. Eisenhauer, E.A., Therasse, P., Bogaerts, J., Schwartz, L.H., Sargent, D., Ford, R., Dancey, J., Arbuck, S., Gwyther, S., Mooney, M., et al. (2009). New response evaluation criteria in solid tumours: revised RECIST guideline (version 1.1). Eur J Cancer 45, 228–247. 10.1016/j.ejca.2008.10.026.

25. Galluzzi, L., Humeau, J., Buqué, A., Zitvogel, L., and Kroemer, G. (2020). Immunostimulation with chemotherapy in the era of immune checkpoint inhibitors. Nat Rev Clin Oncol 17, 725–741. 10.1038/s41571-020-0413-z.

26. Caushi, J.X., Zhang, J., Ji, Z., Vaghasia, A., Zhang, B., Hsiue, E.H., Mog, B.J., Hou, W., Justesen, S., Blosser, R., et al. (2021). Transcriptional programs of neoantigen-specific TIL in anti-PD-1-treated lung cancers. Nature 596, 126–132. 10.1038/s41586-021-03752-4.

27. Lowery, F.J., Krishna, S., Yossef, R., Parikh, N.B., Chatani, P.D., Zacharakis, N., Parkhurst, M.R., Levin, N., Sindiri, S., Sachs, A., et al. (2022). Molecular signatures of antitumor neoantigen-reactive T cells from metastatic human cancers. Science 375, 877–884. 10.1126/science.abl5447.

28. Hanada, K.I., Zhao, C., Gil-Hoyos, R., Gartner, J.J., Chow-Parmer, C., Lowery, F.J., Krishna, S., Prickett, T.D., Kivitz, S., Parkhurst, M.R., et al. (2022). A phenotypic signature that identifies neoantigen-reactive T cells in fresh human lung cancers. Cancer Cell 40, 479–493.e476. 10.1016/j.ccell.2022.03.012.

29. Liu, Z., Yang, Z., Wu, J., Zhang, W., Sun, Y., Zhang, C., Bai, G., Yang, L., Fan, H., Chen, Y., et al. (2025). A single-cell atlas reveals immune heterogeneity in anti-PD-1-treated non-small cell lung cancer. Cell. 10.1016/j.cell.2025.03.018.

30. Taube, J.M., Anders, R.A., Young, G.D., Xu, H., Sharma, R., McMiller, T.L., Chen, S., Klein, A.P., Pardoll, D.M., Topalian, S.L., and Chen, L. (2012). Colocalization of inflammatory response with B7-h1 expression in human melanocytic lesions supports an adaptive resistance mechanism of immune escape. Sci Transl Med 4, 127ra137. 10.1126/scitranslmed.3003689.

31. Velcheti, V., Schalper, K.A., Carvajal, D.E., Anagnostou, V.K., Syrigos, K.N., Sznol, M., Herbst, R.S., Gettinger, S.N., Chen, L., and Rimm, D.L. (2014). Programmed death ligand-1 expression in non-small cell lung cancer. Lab Invest 94, 107–116. 10.1038/labinvest.2013.130.

32. Schad, S.E., Chow, A., Mangarin, L., Pan, H., Zhang, J., Ceglia, N., Caushi, J.X., Malandro, N., Zappasodi, R., Gigoux, M., et al. (2022). Tumor-induced double positive T cells display distinct lineage commitment mechanisms and functions. J Exp Med 219. 10.1084/jem.20212169.

33. Chow, A., Uddin, F.Z., Liu, M., Dobrin, A., Nabet, B.Y., Mangarin, L., Lavin, Y., Rizvi, H., Tischfield, S.E., Quintanal-Villalonga, A., et al. (2023). The ectonucleotidase CD39 identifies tumor-reactive CD8. Immunity 56, 93–106.e106. 10.1016/j.immuni.2022.12.001.

34. Wolfl, M., Kuball, J., Ho, W.Y., Nguyen, H., Manley, T.J., Bleakley, M., and Greenberg, P.D. (2007). Activation-induced expression of CD137 permits detection, isolation, and expansion of the full repertoire of CD8+ T cells responding to antigen without requiring knowledge of epitope specificities. Blood 110, 201–210. 10.1182/blood-2006-11-056168.

35. Game, D.S., Cao, X., and Jiang, S. (2008). Regulatory T cells: the cunning fox and its clinical application. Sci Signal 1, mr3. 10.1126/scisignal.151mr3.

36. Le, D.T., Ladle, B.H., Lee, T., Weiss, V., Yao, X., Leubner, A., Armstrong, T.D., and Jaffee, E.M. (2011). CD8⁺ Foxp3⁺ tumor infiltrating lymphocytes accumulate in the context of an effective anti-tumor response. Int J Cancer 129, 636–647. 10.1002/ijc.25693.

37. Conde, E., Casares, N., Mancheño, U., Elizalde, E., Vercher, E., Capozzi, R., Santamaria, E., Rodriguez-Madoz, J.R., Prosper, F., Lasarte, J.J., et al. (2023). FOXP3 expression diversifies the metabolic capacity and enhances the efficacy of CD8 T cells in adoptive immunotherapy of melanoma. Mol Ther 31, 48–65. 10.1016/j.ymthe.2022.08.017.

38. Churlaud, G., Pitoiset, F., Jebbawi, F., Lorenzon, R., Bellier, B., Rosenzwajg, M., and Klatzmann, D. (2015). Human and Mouse CD8(+)CD25(+)FOXP3(+) Regulatory T Cells at Steady State and during Interleukin-2 Therapy. Front Immunol 6, 171. 10.3389/fimmu.2015.00171.

39. Tassi, E., Grazia, G., Vegetti, C., Bersani, I., Bertolini, G., Molla, A., Baldassari, P., Andriani, F., Roz, L., Sozzi, G., et al. (2017). Early Effector T Lymphocytes Coexpress Multiple Inhibitory Receptors in Primary Non-Small Cell Lung Cancer. Cancer Res 77, 851–861. 10.1158/0008-5472.CAN-16-1387.

40. Heeren, A.M., Rotman, J., Stam, A.G.M., Pocorni, N., Gassama, A.A., Samuels, S., Bleeker, M.C.G., Mom, C.H., Zijlmans, H.J.M.A., Kenter, G.G., et al. (2019). Efficacy of PD-1 blockade in cervical cancer is related to a CD8. J Immunother Cancer 7, 43. 10.1186/s40425-019-0526-z.

41. Walker, C.R., and Angelo, M. (2024). Insights and Opportunity Costs in Applying Spatial Biology to Study the Tumor Microenvironment. Cancer Discov 14, 707–710. 10.1158/2159-8290.CD-24-0348.

42. Backman, M., Strell, C., Lindberg, A., Mattsson, J.S.M., Elfving, H., Brunnström, H., O’Reilly, A., Bosic, M., Gulyas, M., Isaksson, J., et al. (2023). Spatial immunophenotyping of the tumour microenvironment in non-small cell lung cancer. Eur J Cancer 185, 40–52. 10.1016/j.ejca.2023.02.012.

43. Moutafi, M.K., Molero, M., Martinez Morilla, S., Baena, J., Vathiotis, I.A., Gavrielatou, N., Castro-Labrador, L., de Garibay, G.R., Adradas, V., Orive, D., et al. (2022). Spatially resolved proteomic profiling identifies tumor cell CD44 as a biomarker associated with sensitivity to PD-1 axis blockade in advanced non-small-cell lung cancer. J Immunother Cancer 10. 10.1136/jitc-2022-004757.

44. Enfield, K.S.S., Colliver, E., Lee, C., Magness, A., Moore, D.A., Sivakumar, M., Grigoriadis, K., Pich, O., Karasaki, T., Hobson, P.S., et al. (2024). Spatial Architecture of Myeloid and T Cells Orchestrates Immune Evasion and Clinical Outcome in Lung Cancer. Cancer Discov 14, 1018–1047. 10.1158/2159-8290.CD-23-1380.

45. Taube, J.M., Roman, K., Engle, E.L., Wang, C., Ballesteros-Merino, C., Jensen, S.M., McGuire, J., Jiang, M., Coltharp, C., Remeniuk, B., et al. (2021). Multi-institutional TSA-amplified Multiplexed Immunofluorescence Reproducibility Evaluation (MITRE) Study. J Immunother Cancer 9. 10.1136/jitc-2020-002197.

46. Fridman, W.H., Meylan, M., Pupier, G., Calvez, A., Hernandez, I., and Sautès-Fridman, C. (2023). Tertiary lymphoid structures and B cells: An intratumoral immunity cycle. Immunity 56, 2254–2269. 10.1016/j.immuni.2023.08.009.

47. Ghiringhelli, F., Bibeau, F., Greillier, L., Fumet, J.D., Ilie, A., Monville, F., Laugé, C., Catteau, A., Boquet, I., Majdi, A., et al. (2023). Immunoscore immune checkpoint using spatial quantitative analysis of CD8 and PD-L1 markers is predictive of the efficacy of anti-PD1/PD-L1 immunotherapy in non-small cell lung cancer. EBioMedicine 92, 104633. 10.1016/j.ebiom.2023.104633.

48. Galon, J., Angell, H.K., Bedognetti, D., and Marincola, F.M. (2013). The continuum of cancer immunosurveillance: prognostic, predictive, and mechanistic signatures. Immunity 39, 11–26. 10.1016/j.immuni.2013.07.008.

49. Brody, R., Zhang, Y., Ballas, M., Siddiqui, M.K., Gupta, P., Barker, C., Midha, A., and Walker, J. (2017). PD-L1 expression in advanced NSCLC: Insights into risk stratification and treatment selection from a systematic literature review. Lung Cancer 112, 200–215. 10.1016/j.lungcan.2017.08.005.

50. Gao, Q., Wang, S., Chen, X., Cheng, S., Zhang, Z., Li, F., Huang, L., Yang, Y., Zhou, B., Yue, D., et al. (2019). Cancer-cell-secreted CXCL11 promoted CD8. J Immunother Cancer 7, 42. 10.1186/s40425-019-0511-6.

51. Ma, Z., Zhang, W., Dong, B., Xin, Z., Ji, Y., Su, R., Shen, K., Pan, J., Wang, Q., and Xue, W. (2022). Docetaxel remodels prostate cancer immune microenvironment and enhances checkpoint inhibitor-based immunotherapy. Theranostics 12, 4965–4979. 10.7150/thno.73152.

52. Garnett, C.T., Schlom, J., and Hodge, J.W. (2008). Combination of docetaxel and recombinant vaccine enhances T-cell responses and antitumor activity: effects of docetaxel on immune enhancement. Clin Cancer Res 14, 3536–3544. 10.1158/1078-0432.CCR-07-4025.

53. Hu, Z., Chen, K., Huang, H., Zhong, X., Wang, Y., Chen, J., He, X., Shi, D., Zeng, Y., Li, J., et al. (2025). The Spatial Proximity of CD8+ FoxP3+PD-1+ Cells to Tumor Cells: A More Accurate Predictor of Immunotherapy Outcomes in Advanced Non-Small-Cell Lung Cancer. Curr Oncol 32. 10.3390/curroncol32050262.

54. Akinboro, O., Vallejo, J.J., Nakajima, E.C., Ren, Y., Mishra-Kalyani, P.S., Larkins, E.A., Vellanki, P.J., Drezner, N.L., Mathieu, L.N., Donoghue, M.B., et al. (2022-6-1). Outcomes of anti–PD-(L)1 therapy with or without chemotherapy (chemo) for first-line (1L) treatment of advanced non–small cell lung cancer (NSCLC) with PD-L1 score ≥ 50%: FDA pooled analysis. Journal of Clinical Oncology 40. 10.1200/JCO.2022.40.16_suppl.9000.

55. Chen, J.H., Nieman, L.T., Spurrell, M., Jorgji, V., Elmelech, L., Richieri, P., Xu, K.H., Madhu, R., Parikh, M., Zamora, I., et al. (2024). Human lung cancer harbors spatially organized stem-immunity hubs associated with response to immunotherapy. Nat Immunol 25, 644–658. 10.1038/s41590-024-01792-2.

56. Lu, S., Succaria, F., Green, B., Giraldo-Castillo, N., Taube, J. (2020/07/01). 906 Early effector T-cell densities predict response in patients with advanced Merkel cell carcinoma treated with anti-PD-1. Journal of Investigative Dermatology 140. 10.1016/j.jid.2020.03.922.

57. Taghizadeh-Popp, M., Kim, J.W., Lemson, G., Medvedev, D., Raddick, M.J., Szalay, A.S., Thakar, A.R., Booker, J., Chhetri, C., Dobos, L., and Rippin, M. (2020). SciServer: A science platform for astronomy and beyond. Astronomy and Computing 33. 10.1016/j.ascom.2020.100412.

58. Reuss, J.E., Anagnostou, V., Cottrell, T.R., Smith, K.N., Verde, F., Zahurak, M., Lanis, M., Murray, J.C., Chan, H.Y., McCarthy, C., et al. (2020). Neoadjuvant nivolumab plus ipilimumab in resectable non-small cell lung cancer. J Immunother Cancer 8. 10.1136/jitc-2020-001282.

59. Stein, J.E., Lipson, E.J., Cottrell, T.R., Forde, P.M., Anders, R.A., Cimino-Mathews, A., Thompson, E.D., Allaf, M.E., Yarchoan, M., Feliciano, J., et al. (2020). Pan-Tumor Pathologic Scoring of Response to PD-(L)1 Blockade. Clin Cancer Res 26, 545–551. 10.1158/1078-0432.CCR-19-2379.

60. Deutsch, J.S., Cimino-Mathews, A., Thompson, E., Provencio, M., Forde, P.M., Spicer, J., Girard, N., Wang, D., Anders, R.A., Gabrielson, E., et al. (2024). Association between pathologic response and survival after neoadjuvant therapy in lung cancer. Nat Med 30, 218–228. 10.1038/s41591-023-02660-6.

61. Deutsch, J.S., Burton, E., Cimino-Mathews, A., Cottrell, T.R., de Andrea, C.E., Fiset, P.O., Gershenwald, J.E., Long, G.V., Messina, J., Salgado, R.F., et al. (2024). 1943P Updated pan-tumor guidelines for neoadjuvant scoring of pathologic response: A joint SITC and INMC effort. Annals of Oncology 35, S1127–S1128. 10.1016/j.annonc.2024.08.2029.

62. Will, E., Green, B., Qadri, A., Jorquera, A., Soto-Diaz, S., Sunshine, J., Stein Deutsch, J., Warrier, G., Lipson, E., Engle, E., et al. (2024). 693 AstroID: A novel REDCap-based relational database to house biospecimen data. Journal of Investigative Dermatology 144, S121. 10.1016/j.jid.2024.06.709.

63. Giraldo, N.A., Nguyen, P., Engle, E.L., Kaunitz, G.J., Cottrell, T.R., Berry, S., Green, B., Soni, A., Cuda, J.D., Stein, J.E., et al. (2018). Multidimensional, quantitative assessment of PD-1/PD-L1 expression in patients with Merkel cell carcinoma and association with response to pembrolizumab. J Immunother Cancer 6, 99. 10.1186/s40425-018-0404-0.

64. Gritsan, A.V., Roskes, J., Sarica, U., Schulze, M., Xiao, M., and Zhou, Y. (2020). New features in the JHU generator framework: Constraining Higgs boson properties from on-shell and off-shell production. Physical Review D 102. 10.1103/PhysRevD.102.056022.

65. Neyman, J. (1933). On the problem of the most efficient tests of statistical hypotheses. Philos T R Soc Lond 231, 289–337. 10.1098/rsta.1933.0009.

66. Gurobi Optimizer Reference Manual (2024).

67. Butler, A., Hoffman, P., Smibert, P., Papalexi, E., and Satija, R. (2018). Integrating single-cell transcriptomic data across different conditions, technologies, and species. Nat Biotechnol 36, 411-+. 10.1038/nbt.4096.

68. Hao, Y.H., Hao, S., Andersen-Nissen, E., Mauck, W.M., Zheng, S.W., Butler, A., Lee, M.J., Wilk, A.J., Darby, C., Zager, M., et al. (2021). Integrated analysis of multimodal single-cell data. Cell 184, 3573-+. 10.1016/j.cell.2021.04.048.

69. Satija, R., Farrell, J.A., Gennert, D., Schier, A.F., and Regev, A. (2015). Spatial reconstruction of single-cell gene expression data. Nat Biotechnol 33, 495–U206. 10.1038/nbt.3192.

70. Stuart, T., Butler, A., Hoffman, P., Hafemeister, C., Papalexi, E., Mauck, W.M., Hao, Y.H., Stoeckius, M., Smibert, P., and Satija, R. (2019). Comprehensive Integration of Single-Cell Data. Cell 177, 1888-+. 10.1016/j.cell.2019.05.031.

71. Korsunsky, I., Millard, N., Fan, J., Slowikowski, K., Zhang, F., Wei, K., Baglaenko, Y., Brenner, M., Loh, P.R., and Raychaudhuri, S. (2019). Fast, sensitive and accurate integration of single-cell data with Harmony. Nat Methods 16, 1289-+. 10.1038/s41592-019-0619-0.

72. Python 3.11.7 Documentation. (2023). https://docs.python.domainunion.de/release/3.11.7/.

73. SciPy v1.12.0 Manual. (2024). https://docs.scipy.org/doc/scipy/.

74. Mason, S.J., and Graham, N.E. (2002). Areas beneath the relative operating characteristics (ROC) and relative operating levels (ROL) curves: Statistical significance and interpretation. Q J Roy Meteor Soc 128, 2145–2166. 10.1256/003590002320603584.

