## Supplemental Information for "Novel Predictive Spatial Biomarker in Non-Small Cell Lung Carcinoma: The Diversity of Niches Unlocking Treatment Sensitivity (DONUTS)"

### Supplemental Tables

**Table S1**. Clinical details for Cohort 1 (NCT02259621), related to Figure 1

**Table S2.** Clinical details for Cohorts 2 and 4 (NCT01673867), related to Figure 1

**Table S3.** Clinical details for Cohort 3 (second line anti-PD-1, standard of care), related to Figure 1

**Table S4.** Steps for accessing the data in AstroPath, related to Figure 1

**Table S5.** Multiplex IF 6-plex assay staining protocol, related to Figure 1

**Table S6**. Top 30 differential genes in CD8+ FoxP3+ (CD4(-)) subcluster compared to CD8+ stem-like memory cells, related to Figure 3

**Table S7.** Distribution of cell lineages among CD8+FoxP3+ niche and background TME neighbor configurations in neoadjuvant cohort, related to Figures 4-5

### Supplemental Figures

**Figure S1**. Consort diagrams illustrating specimen exclusion criteria for tumor tissue specimens from four NSCLC patient cohorts

**Figure S2**. Quantitatively validated multiplex IF staining demonstrates association of pre-treatment cell densities with patient outcomes in patients receiving anti-PD-1 or chemotherapy, related to Figure 1

**Figure S3**. Single-cell RNA sequencing highlights early, effector phenotype of the CD8+FoxP3+ cells, related to Figure 3

**Figure S4.** The CD8+FoxP3+ cell effector phenotype is identified in on-treatment specimens from patients receiving neoadjuvant anti-PD-1 plus chemotherapy, related to Figure 3

**Figure S5**. Across four independent cohorts, the contact neighbors of CD8+FoxP3+ T-cells are enriched in cytotoxic CD8+FoxP3(-) T-cells and depleted in tumor cells – an immunoactive niche that is recapitulated in the DONUTS, related to Figures 4-5

**Figure S6.** Quantification of the DONUTS stratifies outcomes in patients with NSCLC receiving anti-PD-1 or chemotherapy, related to Figures 4-5

**Figure S7**. Development and validation of the Discriminant of Niches Unlocking Treatment Sensitivity (DONUTS), related to Figures 4-5

**Supplemental Table S1. Clinical details for Cohort 1 (NCT02259621)**

| **Characteristic** - no. (%)* | **Cohort 1** (n=25) | **CD8+FoxP3+ High** (n=19) | **CD8+FoxP3+ Low** (n=6) |
|---|---|---|---|
| **Age at Collection** - years | | | |
| Median (range) | 67 (48-84) | 67 (51-79) | 66 (48-84) |
| **Sex** | | | |
| Female | 11 (44) | 10 (53) | 1 (17) |
| Male | 14 (56) | 9 (47) | 5 (83) |
| **NSCLC Type** | | | |
| Adenocarcinoma | 13 (52) | 11 (58) | 2 (33) |
| Squamous-cell carcinoma | 6 (24) | 3 (16) | 3 (50) |
| Other | 6 (24) | 5 (26) | 1 (17) |
| **Pre-Treatment Disease Stage** | | | |
| Stage I | 5 (20) | 4 (21) | 1 (17) |
| Stage II | 11 (44) | 8 (42) | 3 (50) |
| Stage IIIA | 9 (36) | 7 (37) | 2 (33) |
| **Neoadjuvant Treatment** | | | |
| Anti-PD-1 alone** | 19 (76) | 15 (79) | 4 (67) |
| Anti-PD-1 + anti-CTLA4*** | 6 (24) | 4 (21) | 2 (33) |
| **Status** | | | |
| Alive | 20 (80) | 17 (89) | 3 (50) |
| Deceased | 5 (20) | 2 (11) | 3 (50) |

*Unless otherwise specified; ** Nivolumab or Pembrolizumab; *** Ipilimumab + Nivolumab

**Supplemental Table S2. Clinical details for Cohorts 2 and 4 (NCT01673867)**

| | **Anti-PD-1 arm** (Nivolumab) | | **Chemotherapy arm** (Docetaxel) | |
|---|---|---|---|---|
| **Survival** (months) | **Clinical Trial** (n=292) | **Cohort 2** (n=14) | **Clinical Trial** (n=290) | **Cohort 4** (n=16) |
| **Overall survival** | | | | |
| Median (95% CI) | 12.2 (9.7-15.0) | 14.5 (2.7-51.9) | 9.4 (8.1-10.7) | 10.5 (8.0-19.6) |
| **Progression-free survival** | | | | |
| Median (95% CI) | 2.3 (2.2-3.3) | 4.7 (1.0-26.5) | 4.2 (3.5-4.9) | 7.5 (1.7-9.1) |

**Supplemental Table S3. Clinical details for Cohort 3 (second line anti-PD-1, standard of care)**

| Characteristic | Cohort 3 (n=20) |
|---|---|
| **Age at collection** - years | |
| Median (range) | 63 (47-78) |
| **Sex** - no. (%) | |
| Female | 3 (15.8) |
| Male | 15 (78.9) |
| Unknown | 1 (5.3) |
| **Second line treatment** - no. (%) | |
| Nivolumab | 15 (78.9) |
| Pembrolizumab | 3 (15.8) |
| Durvalumab | 1 (5.3) |
| **Best overall radiographic response** - no. (%) | |
| Progressive disease | 13 (65%) |
| Partial response or stable disease | 6 (30%) |
| Unknown | 1 (5%) |

**Table S4. Steps for accessing the data in AstroPath**

| |
|---|
| Instructions for data access are here: https://www.sciserver.org/integration/astropath/ |
| Start by creating a SciServer account and log in on https://www.sciserver.org/. |
| Next you will need to join the AstroPath science domain: |
| 1. navigate to https://apps.sciserver.org/dashboard/science/ |
| 2. select AstroPath from the left menu |
| 3. click the green "Join" button |

### Supplemental Table S5. Multiplex IF 6-plex assay staining protocol

| Marker | Primary Antibody | | | | Secondary Antibody | | | TSA | |
|---|---|---|---|---|---|---|---|---|---|
| | Species | Clone | Source | Final [µg/mL] | Amplification | Source | Final [µg/mL] | Opal | Dilution |
| FoxP3 | Mouse | 236A/E7 | Abcam | 5 | PowerVision poly-HRP anti-Mouse IgG | PV6114, Leica | 1to2** | 570 | 1to200 |
| CD8 | Mouse | 4B11 | AbD Serotic | 1to100* | 1X Opal Anti-Ms + Rb HRP | ARH1001EA, Akoya | RTU | 540 | 1to100 |
| Cytokeratin | Mouse | AE1/AE3 | DAKO | 0.876 | 1X Opal Anti-Ms + Rb HRP | ARH1001EA, Akoya | RTU | 620 | 1to100 |
| PD-1 | Rabbit | EPR4877(2) | Abcam | 0.25 | PowerVision poly-HRP anti-Rabbit IgG | PV6119, Leica | 1to2** | 650 | 1to100 |
| PD-L1 | Rabbit | SP142 | Spring Bioscience | 0.2 | PowerVision poly-HRP anti-Rabbit IgG | PV6119, Leica | 1to2** | 520 | 1to100 |
| CD163 | Mouse | 10D6 | Abcam | 0.49 | 1X Opal Anti-Ms + Rb HRP | ARH1001EA, Akoya | RTU | 690 | 1to50 |

*RTU Anti-CD8 was diluted 1:100. All antibodies were diluted in Antibody Diluent Background Reducing (S3022, Dako).
**RTU PowerVision poly-HRP anti-Mouse IgG and PowerVision poly-HRP anti-Rabbit IgG were diluted 1:2. All antibodies were diluted in 1X PBS.

**Supplemental Table S6. Top 30 differential genes in CD8+CD4-FoxP3+ subcluster compared to CD8+ stem-like memory cells.**

| Gene | Avg_log2FC | Adjusted p-val |
|---|---|---|
| FOXP3 | 3.19639808 | 0 |
| GZMH | 2.23586945 | 8.99E-138 |
| CCL4 | 2.43349407 | 2.15E-115 |
| GZMB | 2.5663902 | 3.51E-114 |
| MIR4435-2HG | 1.59139198 | 1.21E-97 |
| CXCL13 | 3.50239444 | 6.20E-95 |
| HLA-DRB1 | 1.79096092 | 2.23E-94 |
| CXCR6 | 1.58792012 | 8.34E-92 |
| CTLA4 | 1.71751134 | 1.01E-88 |
| DUSP4 | 1.49891419 | 6.22E-82 |
| TNFRSF9 | 1.58043987 | 1.09E-81 |
| HLA-DQA1 | 1.71426225 | 6.33E-81 |
| HLA-DRA | 1.79262425 | 7.68E-81 |
| CCL4L2 | 2.65935797 | 2.19E-71 |
| GZMA | 1.62550411 | 3.24E-71 |
| TIGIT | 1.43046169 | 2.85E-61 |
| ENTPD1 | 1.06456219 | 3.62E-61 |
| BATF | 1.64687986 | 3.76E-61 |
| CARD16 | 1.43390206 | 1.21E-58 |
| VCAM1 | 1.28842847 | 3.10E-57 |
| AC243829.4 | 0.8312109 | 3.65E-52 |
| CD4 | 0.88000281 | 4.55E-50 |
| SLA2 | 1.25242457 | 6.89E-49 |
| CYTOR | 1.15329948 | 4.11E-48 |
| CCL3 | 1.51592846 | 1.36E-47 |
| RGS1 | 1.19417745 | 1.57E-46 |
| HLA-DRB5 | 1.18867832 | 3.06E-45 |
| KRT86 | 1.52449593 | 3.26E-43 |
| CCR8 | 0.56135392 | 8.30E-42 |
| PRF1 | 1.20927982 | 6.04E-40 |

**Supplemental Table S7. Distribution of cell lineages among the CD8+FoxP3+ niche and background TME niches in Cohort 1.**

| | CD8+FoxP3+ niche (D≥0.45) | | Background TME (D<0.45) | |
|---|---|---|---|---|
| | **Training samples** (n=6) | **Analysis samples** (n=25) | **Training samples** (n=6) | **Analysis samples** (n=25) |
| **Other** | 1,855,852 | 737,893 | 5,656,561 | 1,595,098 |
| **CD8+** | 1,004,621 | 227,294 | 636,544 | 54,295 |
| **FoxP3+** | 264,159 | 128,352 | 304,741 | 77,826 |
| **Tumor** | 84,155 | 34,559 | 1,699,878 | 841,430 |
| **CD163+** | 432,240 | 182,300 | 802,098 | 289,211 |
| **CD8+FoxP3+** | 15,255 | 5,542 | 34,508 | 6,224 |

**Supplemental Figure S1.**

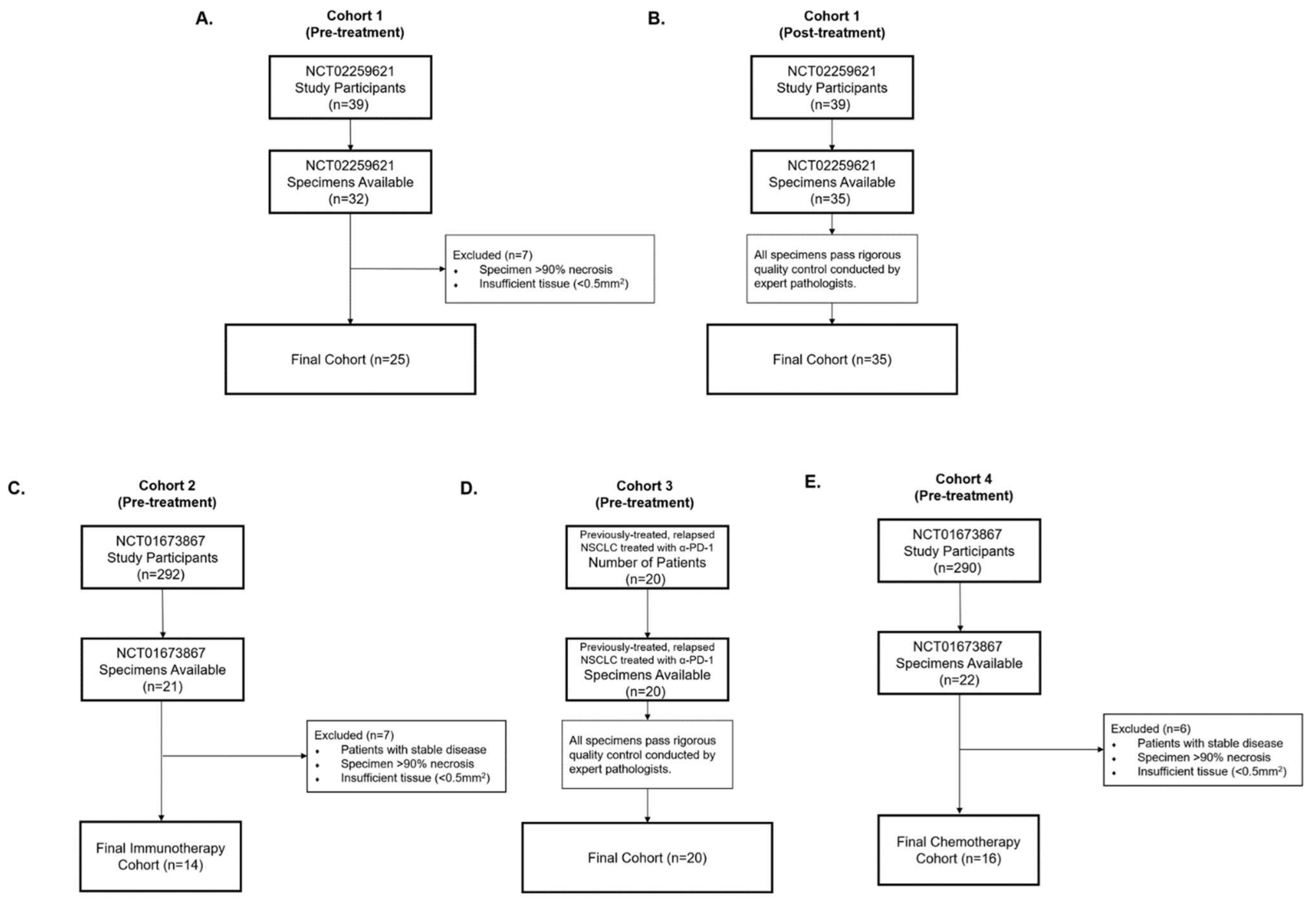


**Supplemental Figure S1. Consort diagrams illustrating specimen exclusion criteria for tumor tissue specimens from four NSCLC patient cohorts.** Patients in cohort one had early stage NSCLC and received neoadjuvant anti-PD-1 based therapy (NCT02259621). **(A)** Available pre-treatment biopsy specimens and **(B)** post-treatment surgical resection specimens were studied with mIF, including n=23 patients with paired pre- and post-treatment specimens. **(C)** Patients in cohort two had advanced NSCLC and received anti-PD-1 based therapy (n=14) in a clinical trial (NCT01673867). **(D)** Patients in cohort three had advanced NSCLC and received anti-PD-1 based therapy following progression on chemotherapy as standard of care. **(E)** Patients in cohort four had advanced NSCLC and received docetaxel chemotherapy (n=16) in a clinical trial (NCT01673867).

**Supplemental Figure S2.**

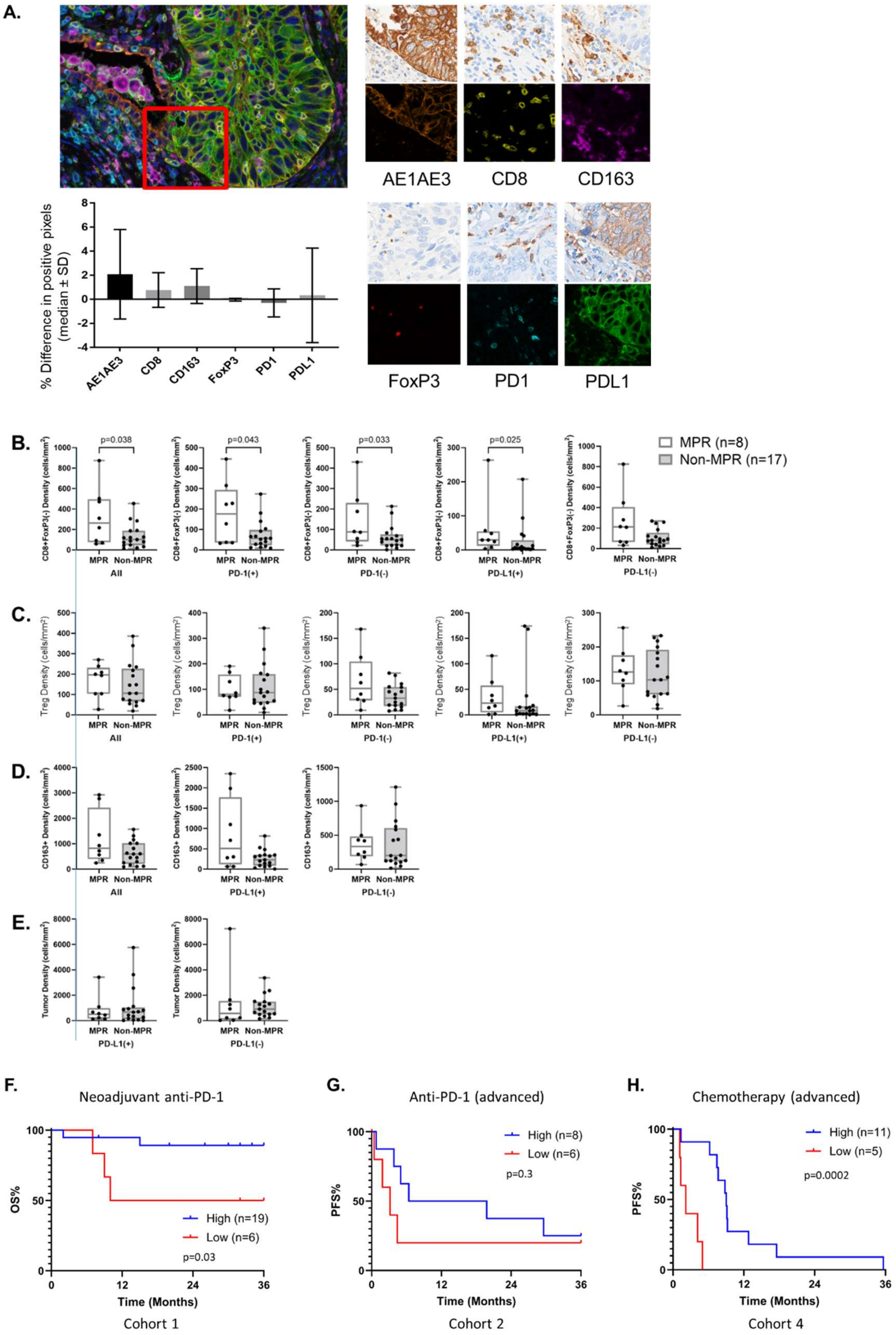

**Supplemental Figure S2. Quantitatively validated multiplex IF staining demonstrates association of pre-treatment cell densities with patient outcomes in patients receiving anti-PD-1 or chemotherapy.**

**(A)** Representative photomicrographs of adjacent serial NSCLC slides stained with multiplex IF and single marker chromogenic IHC. For each marker, the percentage of positive pixels was quantified for both mIF and IHC staining. Less than 5% difference in positive pixels was demonstrated between mIF and IHC staining across N=8 archival NSCLC resection specimens (bottom left plot, n=4 each of adenocarcinomas and squamous cell carcinomas). **(B-E)** Comparison of pre-treatment non-CD8+FoxP3+ cell subset densities between patients whose definitive resection specimen showed MPR vs. non-MPR (Cohort 1). For each cellular subset, densities are presented for all cells, shown also by PD-1 and/or PD-L1 status. All CD8+FoxP3(-) cell subsets except PD-L1(-) were significant higher in pre-treatment biopsies of tumors with MPR following anti-PD-1 versus non-MPR. **(F)** Overall survival (OS) of patients with resectable NSCLC receiving neoadjuvant anti-PD-1 (Cohort 1) stratified by pre-treatment CD8+FoxP3+ T-cell densities. **(G)** Progression free survival (PFS) of patients with advanced pre-treated NSCLC receiving anti-PD-1 (Cohort 2) stratified by pre-treatment CD8+FoxP3+ T-cell densities. **(H)** PFS in patients with advanced pre-treated NSCLC receiving second-line chemotherapy (Cohort 4) stratified by pre-treatment CD8+FoxP3+ T-cell densities.

**Supplemental Figure S3.**

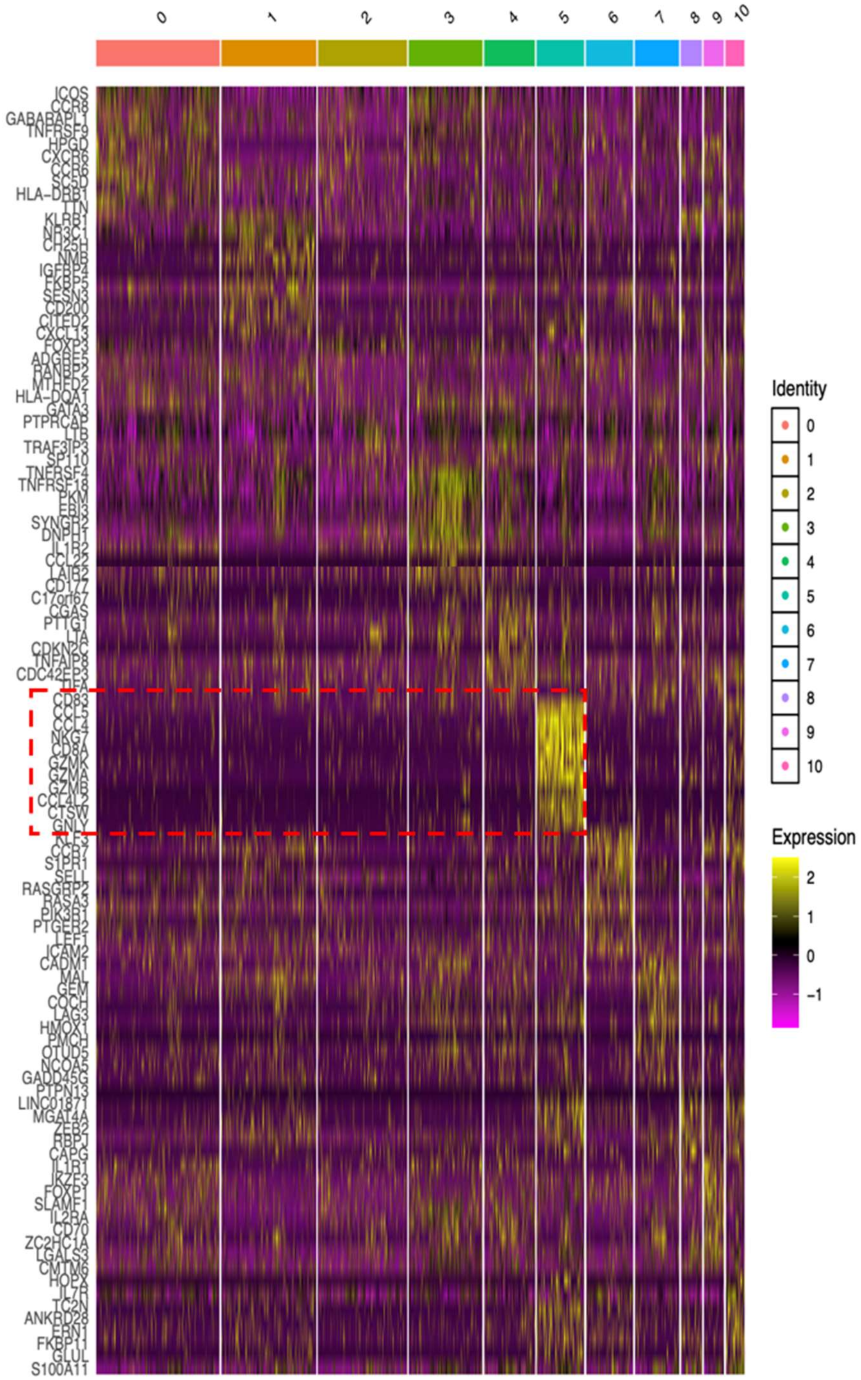


**Supplemental Figure S3. Single-cell RNA sequencing highlights early, effector phenotype of the CD8+FoxP3+ cells.** A heatmap shows the gene expression profiles of 17,210 FoxP3+ TILs that passed quality control in 11 UMAP-derived unique cell clusters. The top 10 differentially expressed genes for each cluster are shown. Cell cluster 5 highlights the early, effector phenotype of the CD8+FoxP3+ T-cells, including coexpression of genes related to cytotoxicity, including granzymes K and B.

**Supplemental Figure S4.**

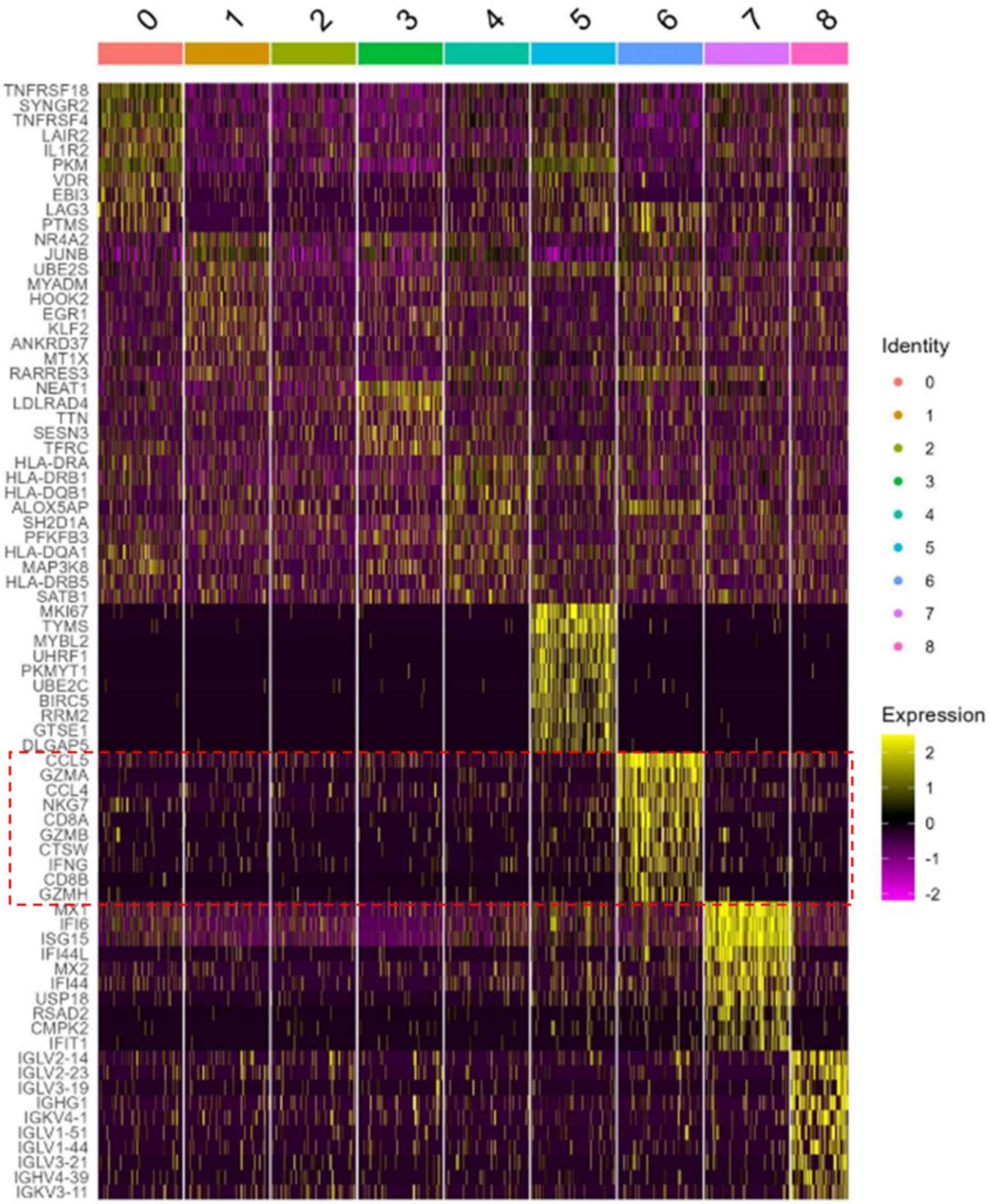


**Supplemental Figure S4. The CD8+FoxP3+ cell effector phenotype is identified in on-treatment specimens from patients receiving neoadjuvant anti-PD-1 plus chemotherapy.** A heatmap shows the gene expression profiles of 31,329 FoxP3+ TILs that passed quality control in 9 UMAP-derived unique cell clusters. The top 10 differentially expressed genes for each cluster are shown. Cell cluster 6 highlights the early, effector phenotype of the CD8+FoxP3+ T-cells, including coexpression of genes related to cytotoxicity, including granzymes K and B.

**Supplemental Figure S5.**

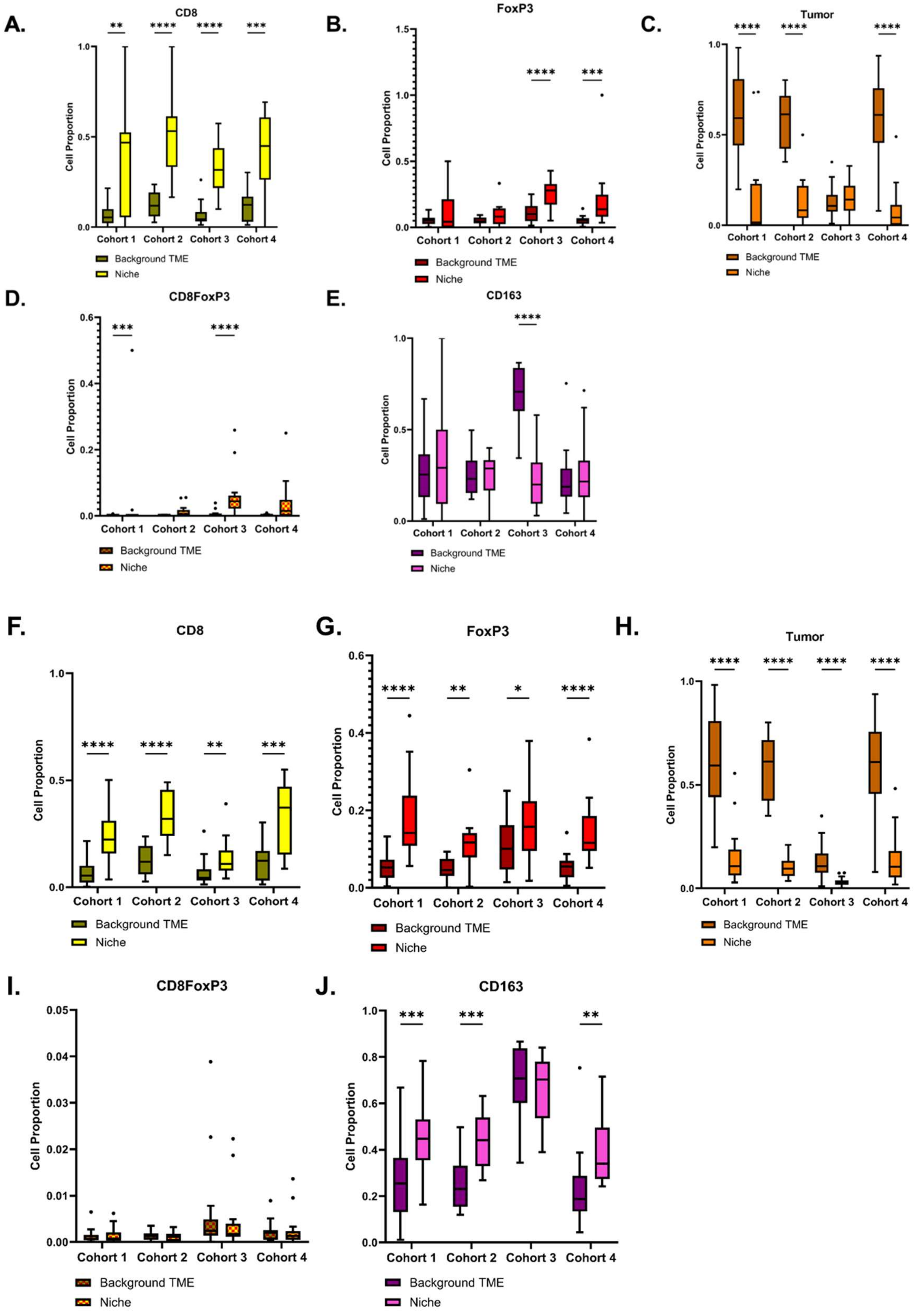

**Supplemental Figure S5. Across four independent cohorts, the contact neighbors of CD8+FoxP3+ T-cells are enriched in cytotoxic CD8+FoxP3(-) T-cells and depleted in tumor cells – an immunoactive niche that is recapitulated in the DONUTS. (A-E)** Boxplots comparing the proportions of each cell lineage in the background TME (dark bars) vs. among CD8+FoxP3+ cell contact neighbors (light bars), including **(A)** cytotoxic (CD8+) T cells, **(B)** regulatory T cells (FoxP3+CD8(-), previously shown to represent CD4+ regulatory T cells (1)), **(C)** tumor cells, **(D)** CD8+FoxP3+ T-cells, and **(E)** macrophages (CD163+). **(F-J)** Boxplots comparing the proportions of each cell lineage in the background TME (dark bars) vs. within the DONUTS, including **(F)** cytotoxic CD8+ T cells, **(G)** regulatory T cells, **(H)** tumor cells, **(I)** CD8+FoxP3+ T-cells, and **(J)** macrophages (CD163+).

**Supplemental Figure S6.**

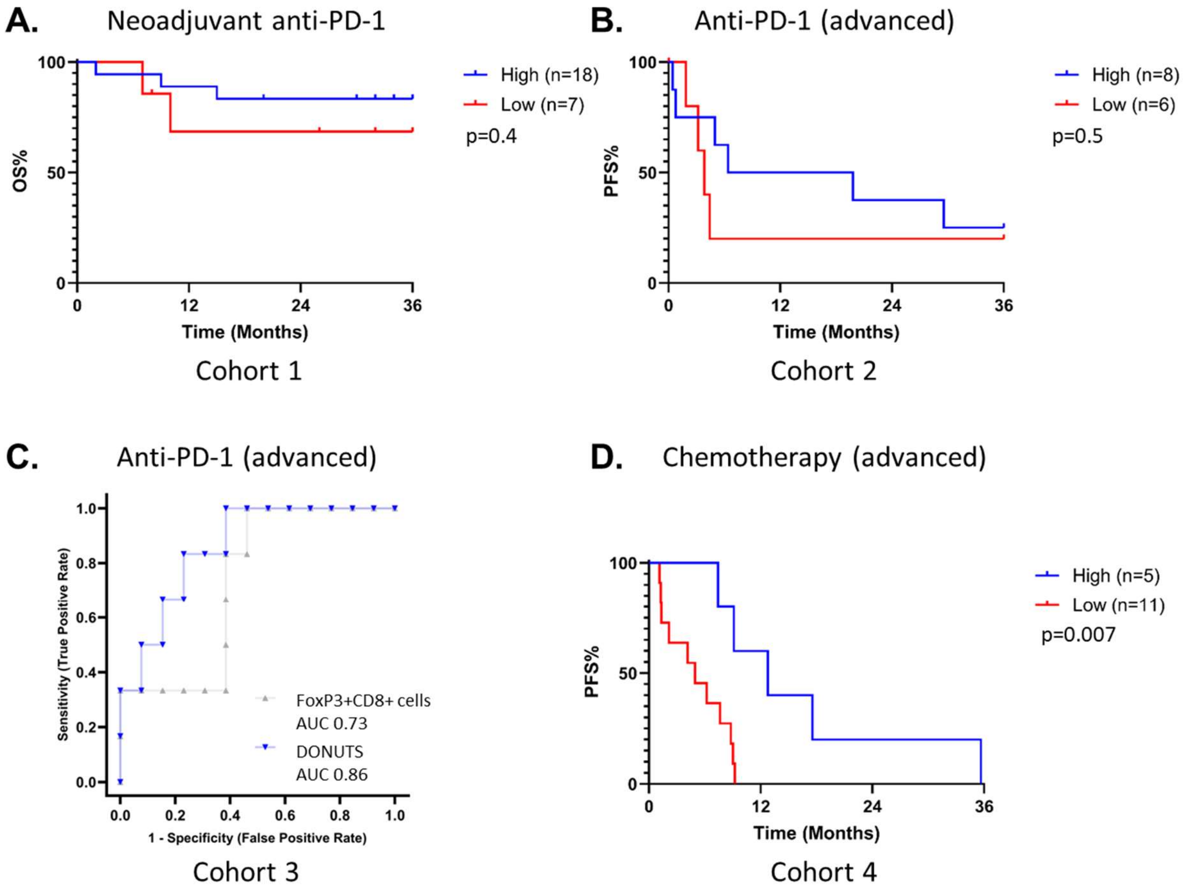


**Supplemental Figure S6. Quantification of the DONUTS stratifies outcomes in patients with NSCLC receiving anti-PD-1 or chemotherapy. (A)** Kaplan Meyer plot showing overall survival (OS) of patients with resectable NSCLC receiving neoadjuvant anti-PD-1 (Cohort 1). **(B)** Kaplan Meyer plot showing progression free survival (PFS) of patients with advanced pre-treated NSCLC receiving anti-PD-1 (Cohort 2). **(C)** ROC curves showing the association of the densities of the DONUTS (blue) and FoxP3+CD8+ cells (gray) with radiographic response in patients with advanced NSCLC treated with anti-PD-1 (Cohort 3). Survival data was not avaiable for Cohort 3. **(D)** Kaplan Meyer plot showing progression free survival (PFS) of patients with advanced pre-treated NSCLC receiving chemotherapy (Cohort 4).

**Supplemental Figure S7**

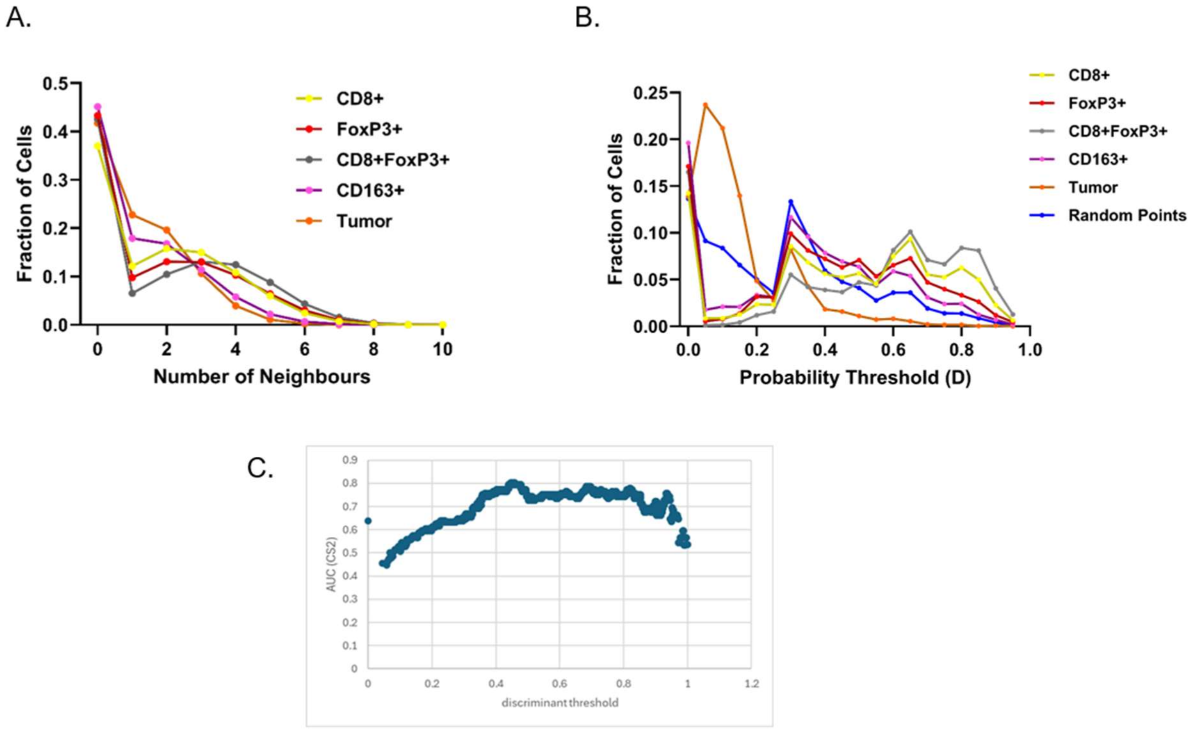


**Supplemental Figure S7. Development and validation of the Diversity of Niches Unlocking Treatment Sensitivity (DONUTS). (A)** Contact neighbors were identified at a distance of 5-10 μm from each cell and there were no differences observed for the number of neighboring cells based on cell lineage (or cell size). (**B**) At increasing thresholds for the likelihood score, the proportion of niches centered on CD8+FoxP3+ T-cells increases (gray) and the proportion of niches centered on other cell lineages or random spatial points decreases. (**C**) AUC values for the association of pre-treatment signal-like niche densities with MPR are shown across all potential likelihood score thresholds. The optimal threshold of t ≥ 0.45 was selected.
